# Dual Targeting of Histone Deacetylases and MYC as Potential Treatment Strategy for H3-K27M Pediatric Gliomas

**DOI:** 10.1101/2024.02.05.578974

**Authors:** Danielle Algranati, Roni Oren, Bareket Dassa, Liat Fellus-Alyagor, Alexander Plotnikov, Haim Barr, Alon Harmelin, Nir London, Guy Ron, Noa Furth, Efrat Shema

## Abstract

Diffuse midline gliomas (DMG) are aggressive and fatal pediatric tumors of the central nervous system that are highly resistant to treatments. Lysine to methionine substitution of residue 27 on histone H3 (H3-K27M) is a driver mutation in DMGs, reshaping the epigenetic landscape of these cells to promote tumorigenesis. H3-K27M gliomas are characterized by deregulation of histone acetylation and methylation pathways, as well as the oncogenic MYC pathway. In search of effective treatment, we examined the therapeutic potential of dual targeting of histone deacetylases (HDACs) and MYC in these tumors. Treatment of H3-K27M patient-derived cells with Sulfopin, an inhibitor shown to block MYC-driven tumors *in-vivo*, in combination with the HDAC inhibitor Vorinostat, resulted in substantial decrease in cell viability. Moreover, transcriptome and epigenome profiling revealed synergistic effect of this drug combination in downregulation of prominent oncogenic pathways such as mTOR. Finally, *in-vivo* studies of patient-derived orthotopic xenograft models showed significant tumor growth reduction in mice treated with the drug combination. These results highlight the combined treatment with PIN1 and HDAC inhibitors as a promising therapeutic approach for these aggressive tumors.

## Introduction

Diffuse midline gliomas (DMGs) are fatal tumors of the central nervous system that occur primarily in children^1,2^. Years of clinical trials have revealed that conventional chemotherapy is ineffective, and with no option for complete resection, DMG is now the leading cause of brain tumor-related death in children^3,4^. Sequencing studies showed that up to 80% of the diagnosed children exhibit a lysine 27 to methionine (H3-K27M) mutation in one of the genes encoding histone H3.3 and H3.1 (clinically classified as DMG, H3 K27M–mutant). Numerous subsequent studies revealed pronounced alterations in the epigenetic landscape driven by this histone mutation^5–8^. Among these global epigenetic alterations are a drastic loss of H3 lysine 27 tri-methylation (H3K27me3), concomitant with an increase in H3 lysine 27 acetylation (H3K27ac). These epigenetic alterations were shown to affect transcriptional programs and support tumorigenesis ^9–14^.

Detailed characterization of the modes of chromatin deregulation in these tumors pointed toward targeting epigenetic pathways as a promising therapeutic approach^15^. Specifically, pervasive H3K27 acetylation patterns observed in H3-K27M DMGs, along with enrichment of H3K27ac specifically on mutant nucleosomes^11–13,16^, underscored the histone acetylation pathway as a potential drug target for this cancer. Indeed, pan-histone deacetylase inhibitors (HDACi), such as Vorinostat and Panobinostat, were found to inhibit growth and to restore gene expression alterations observed in H3-K27M malignant gliomas^17–19^. However, recent studies demonstrated that H3-K27M cells develop resistance to HDACi treatment^17,19^, stressing the importance of identifying novel drug combinations to induce synergistic inhibition of oncogenic pathways.

The MYC oncogenic pathway is frequently deregulated in diverse cancers, and thus stands as a prominent therapeutic target^20^. High-level gene amplification of *MYC* is recurrently observed in pediatric high-grade gliomas^21,22^. H3-K27M gliomas show high expression of *MYC* and MYC target genes, due to both epigenetic alterations and structural variants, resulting in a viability dependency on MYC signaling^12,23,24^. Unfortunately, directly targeting MYC in tumors is extremely challanging^25^. Dubiella et al. have recently developed a unique inhibitor targeting Peptidyl-prolyl isomerase NIMA-interacting-1 (PIN1), an upstream regulator of MYC activation^26^. PIN1 itself is frequently upregulated in many types of cancers, resulting in sustained proliferative signaling and tumor growth^27,28^. The PIN1 inhibitor, Sulfopin, was shown to downregulate MYC target genes *in-vitro* and to block MYC-driven tumors in murine models of neuroblastoma^26^.

To address the increasing need for an effective combination therapy for H3-K27M DMGs, we tested whether dual targeting of histone acetylation and MYC activation, using Vorinostat and Sulfopin, would yield beneficial outcome. We show that the combined treatment substantially reduces viability of patient-derived glioma cells, in concordance with inhibition of prominent oncogenic pathways, including the mTOR pathway. Treatment of H3-K27M DMG xenograft mice models with this drug combination led to reduced tumor growth and confirmed downregulation of mTOR *in-vivo*. Together, these results suggest the combination of PIN1 and HDAC inhibitors as a promising therapeutic strategy for H3-K27M gliomas.

## Results

### Sulfopin inhibits MYC targets and reduces viability of H3-K27M glioma cells

In light of the important role of MYC in H3-K27M DMGs, we set to explore whether the PIN1 inhibitor, Sulfopin, may affect MYC signaling in these cells and thus confer therapeutic potential. Sulfopin dependent inhibition of PIN1 was shown to downregulate the expression of MYC target genes in several cancer cell lines^26^. Indeed, RNA sequencing analysis of H3-K27M mutant patient-derived glioma cells (SU-DIPG13) treated with Sulfopin revealed strong enrichment of MYC targets among the downregulated genes (Fig. 1A). The genes downregulated by Sulfopin were also enriched for MYC-bound genes identified in several non-glioma cell lines, supporting a role for PIN1 in activation of the MYC pathway also in these glioma cells (Fig. S1A-B). Of note, Sulfopin was developed as a covalent inhibitor of PIN1, and did not lead to PIN1 degradation, as previously shown (Fig. S1C) ^26^. We further validated the decreased expression of prominent MYC targets^27,29,30^ upon Sulfopin treatment by RT-qPCR analysis (Fig. 1B, S1D). Interestingly, Cut&Run analysis^31^ revealed an increase in H3K27me3 levels, indicative of repressed chromatin^32^, around the transcriptional start sites (TSS) of MYC target genes, correlating with silencing of these genes upon PIN1 inhibition (Fig. 1C, S1E). Of note, global H3K27me3 distribution on all TSS and gene-body regions was not affected by Sulfopin treatment, supporting a specific increase in H3K27me3 on the TSS of MYC target genes (Fig. S1F-G). Furthermore, Sulfopin-unique H3K27me3 peaks (i.e., peaks that were found only in the Sulfopin-treated cells, and not in DMSO-treated cells) were enriched for MYC target genes (Fig. 1D).

**Figure 1:**
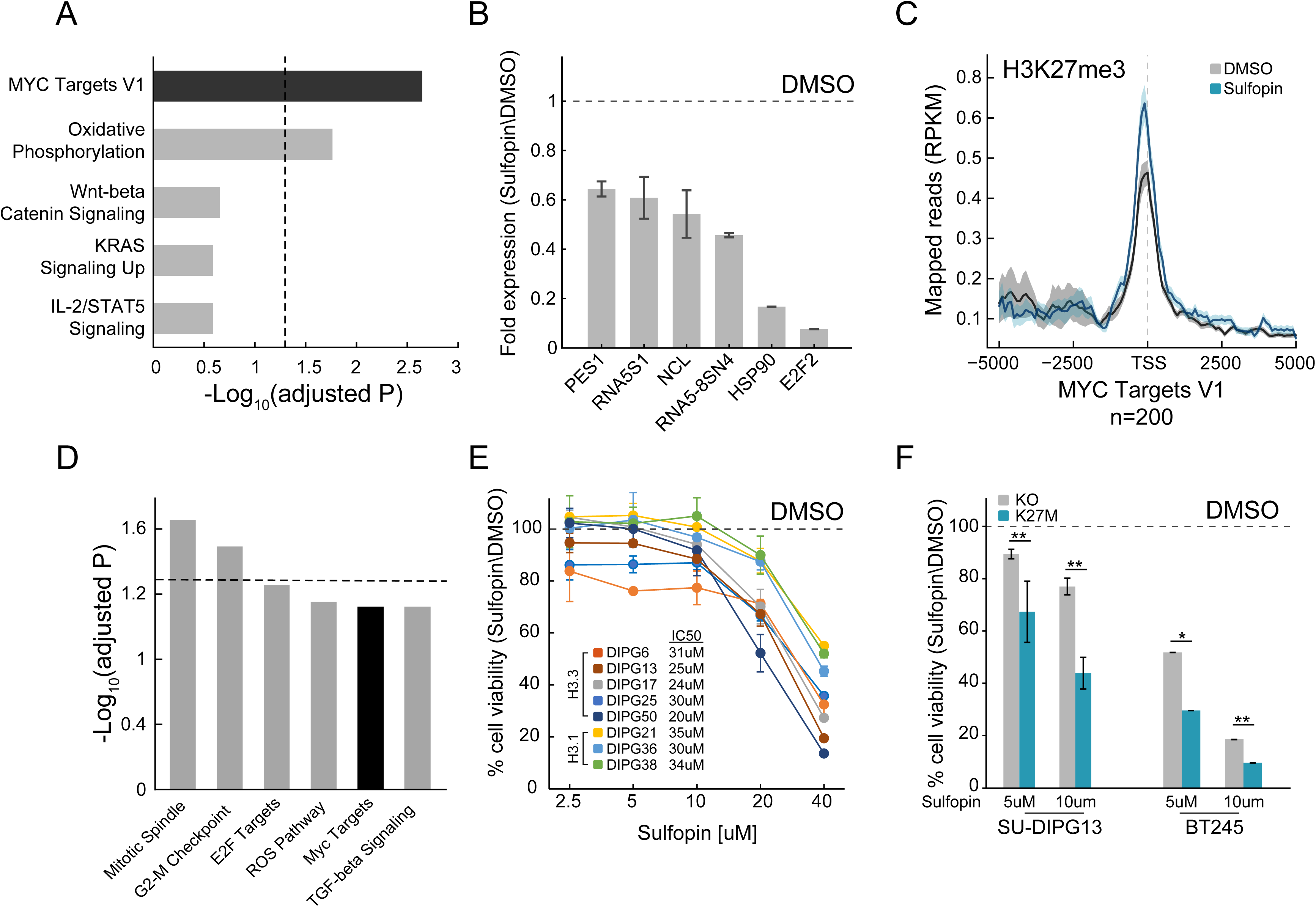
Sulfopin inhibits MYC signaling and reduces viability of DMG cells in a H3-K27M-dependent manner. (A) Functional enrichment analysis on significantly downregulated genes in SU-DIPG13 cells treated with 10uM Sulfopin for 12 hours, compared to DMSO. Enrichr algorithm^123^ was used to compare downregulated genes against the Molecular Signatures Database (MSigDB) hallmark geneset^119^. Dashed line denotes adjusted p-value = 0.05. MYC targets are significantly enriched among the Sulfopin downregulated genes. (B) RT-qPCR analysis of selected MYC target genes in SU-DIPG13 cells treated with 10uM Sulfopin for 12 hours, compared to DMSO. Fold change between Sulfopin and DMSO treated cells was calculated and the mean ± SD of two technical repeats is shown. (C) H3K27me3 Cut&Run read coverage over MYC target genes (‘MYC Targets V1’ hallmark geneset^119^, n=200), in SU-DIPG13 cells treated with 10uM Sulfopin for 8 days compared to DMSO. Sulfopin treatment increases H3K27me3 levels on the TSS of MYC target genes. (D) Functional enrichment analysis of the genes associated with Sulfopin-unique H3K27me3 peaks, in SU-DIPG13 cells treated as in C. Dashed line denotes adjusted p-value = 0.05. MYC target geneset (‘MYC Targets V1’ hallmark geneset^119^) is mildly enriched among these genes, with adjusted p-value of 0.077. (E) Cell viability, as measured by CellTiterGlo, of eight DMG cultures (H3.3K27M: SU-DIPG13, SU-DIPG6, SU-DIPG17, SU-DIPG25 and SU-DIPG50. H3.1K27M: SU-DIPG36, SU-DIPG38 and SU-DIPG21), treated with Sulfopin for eight days with pulse at day four, compared to DMSO. Mean±SD of two technical replicates is shown. Logarithmic scale is used for the x-axis. Sulfopin treatment led to a mild reduction in cell viability in all H3-K27M glioma cultures. (F) Cell viability, as measured by CellTiterGlo, of two isogenic DMG cell lines (SU-DIPG13 and BT245) in which the mutant histone was knocked-out (KO), treated with the indicated concentration of Sulfopin for 8 days, compared to DMSO. For each cell line and concentration, the fold change in viability between Sulfopin and DMSO treated cells is shown. For SU-DIPG13-mean ± SE of at least two independent experiments is shown. For BT245-mean± SD of three technical replicates is shown. H3-K27M glioma cells show higher sensitivity to Sulfopin treatment compared to the KO cells. *P < 0.05; **P < 0.01 (two-sample t-test over all technical replicates). Significance adjusted after Bonferroni correction.

We next explored the effect of Sulfopin on the viability of eight patient derived DMG cultures, harboring either the H3.3-K27M or H3.1-K27M mutation. Inhibition of PIN1 by Sulfopin was previously shown to delay cell cycle progression and thus reduce cell viability after extended treatments of 6-8 days ^26^. In line with downregulation of MYC signaling, Sulfopin treatment led to a mild reduction in cell viability, conserved across the different cultures (Fig. 1E). To examine whether H3-K27M mutation renders DMG cells more sensitive to Sulfopin treatment, we took advantage of two sets of isogenic H3-K27M mutant patient-derived DMG cell-lines (SU-DIPG13, BT245) in which the *H3F3A*-K27M mutant allele has been knocked-out (KO). In both lines, the reduction in cell viability was partially rescued in the KO cells (Fig. 1F), suggesting higher sensitivity of H3-K27M mutant cells to MYC inhibition by Sulfopin. These results are in line with previous reports showing H3-K27M-dependent activation of the MYC/RAS axis in mouse and human cells^12,23,24^. Of note, DMG cells showed greater sensitivity to the inhibitor compared to human astrocytic-like cells (Fig. S1H).

To further explore the status of MYC and PIN1 as indicators of response to Sulfopin inhibition, we analyzed published transcriptomic dataset of DMG tumors^35^. Interestingly, while DMG samples showed significantly higher expression of MYC and its target genes compared to normal samples, PIN1 expression was lower in the cancer compared to matched normal tissue (Fig. S1I-K). Furthermore, H3-K27M mutant tumors showed higher expression of MYC, compared to H3 WT tumors, while PIN1 levels were reduced across samples^22^ (Fig. S1L-M). These results suggest that MYC expression levels, and not PIN1, may determine the sensitivity of H3-K27M cells to Sulfopin treatment. To examine this notion, we measured the expression levels of MYC, as well as the doubling time of the different DMG cultures. The response to Sulfopin treatment (as reflected by IC50 measurements) correlated with the cells’ doubling time (Fig. S1N). Cells that divide faster were more sensitive to Sulfopin treatment, in line with PIN1 role in regulating cell cycle progression, and specifically G2/M transition^36,37^. As expected, the doubling time negatively correlated with MYC expression, suggesting that MYC levels may predict the response to Sulfopin (Fig S1O). Indeed, MYC levels negatively correlated with the IC50 of Sulfopin; thus, cells relaying on high MYC activity showed higher sensitivity to Sulfopin (Fig S1P).

H3K27me3 levels were shown to be drastically deregulated in DMG cells expressing H3-K27M, with global loss of this modification across the genome, and retention of residual H3K27me3 at CpG islands (CGIs). These residual H3K27me3 levels were shown to be necessary to silence tumor suppressor genes and maintain cell proliferation in DMG cells^33,34^. Interestingly, Sulfopin treatment resulted in reduction of H3K27me3 signal over CGIs, perhaps contributing to Sulfopin-dependent growth inhibition in these cells (Fig. S1Q). Taken together, our results indicate that PIN1 inhibition by Sulfopin downregulated MYC target genes, reduced H3K27me3 levels at CGIs, and led to reduce viability of DMG cells in a H3-K27M-dependant manner.

### Combination of Sulfopin and Vorinostat elicits robust inhibition of oncogenic pathways and an additive effect on cell viability

Inspired by the positive outcome of combinatorial drug therapy in H3-K27M DMG models^17,38^, we aimed to explore whether combining Sulfopin treatment with an epigenetic drug will further reduce cell viability. H3-K27M gliomas are characterized by deregulation of epigenetic pathways affecting post-translational modifications of lysine 27 (methylation and acetylation) and lysine 4 (methylation) on histone H3^9–12,16,39^. Thus, we explored the effects of combining sub-IC50 concentration of Sulfopin with four epigenetic inhibitors targeting critical components associated with these modifications: (1) EPZ-6438, an inhibitor of EZH2, the catalytic subunit of the PRC2 complex that deposits H3K27me3^40^. H3-K27M cells were reported to be sensitive to EZH2 inhibition as they rely on the residual H3K27me3 levels to suppress key tumor-suppressor genes^13,33,41,42^; (2) GSK-J4, an inhibitor of the H3K27me3 demethylase JMJD3^43^, shown to rescue H3K27me3 levels in H3-K27M gliomas^17,44^; (3) MM-102, an inhibitor of the H3K4me3 methyltransferase MLL1^40^, shown to reduce viability of H3-K27M-mutant cells^16^; and (4) Vorinostat, an HDAC inhibitor that was shown to have clinical benefit in H3-K27M gliomas^17–19^. Treatment dosages of these drugs were set according to previous studies ^33,17,16^. We found that the combination of Sulfopin and Vorinostat (HDACi) had the strongest effect, reducing viability by 80% (Fig. S2A).

Next, we adjusted treatment protocols to account for the optimal treatment period for each drug, and examined the effect of this drug combination on the viability of eight H3-K27M mutant patient-derived DMG cultures, across different dosages (Fig. 2A-B and S2B-D). For each pair of concentrations, we calculated a BLISS index; an additive effect of the two drugs was determined as BLISS index ∼1^45,46^ (Fig. 2C, S2E). While the combined treatment was effective in all the cultures, cells harboring H3.3-K27M showed higher sensitivity (Fig. 2D, S2F). Moreover, while an additive effect of Sulfopin and Vorinostat was detected in all the H3.3-K27M DMG cultures in most of the drug doses, for the H3.1-K27M cultures we measured an additive effect only in higher dosages of Vorinostat (Fig. 2C-E and S2E-F). In line with the difference in sensitivity, H3.1- and H3.3-K27M cells differed substantially in MYC expression levels, with H3.1-K27M cells expressing significantly lower levels of MYC (Fig. S2G). Indeed, BLISS indexes negatively correlated with MYC expression levels (Fig. 2F), indicating that H3.3-K27M cells, which express higher levels of MYC, are more sensitive to the combination of Sulfopin and Vorinostat. Importantly, this highlights MYC, and MYC target genes, as potential biomarkers of response for the combined treatment. For example, the expression levels of the two MYC targets, NOC4L and PSMD3, negatively correlated with the BLISS indexes and significantly differed between H3.1- and H3.3-K27M cells (Fig. 2F and S2H). Furthermore, these and additional MYC targets were downregulated following the combined treatment in the H3.3-K27M culture, SU-DIPG13 (Fig. S2I). Overall, our data demonstrates that combining Sulfopin treatment with Vorinostat results in an additive effect on the viability of DMG cultures, with H3.3-K27M cells showing higher sensitivity. This is in line with previous studies highlighting clinical and molecular distinction between H3.1- and H3.3-K27M mutant tumors, which is thought to dictate their targetable vulnerabilities^5,47,48^.

**Figure 2:**
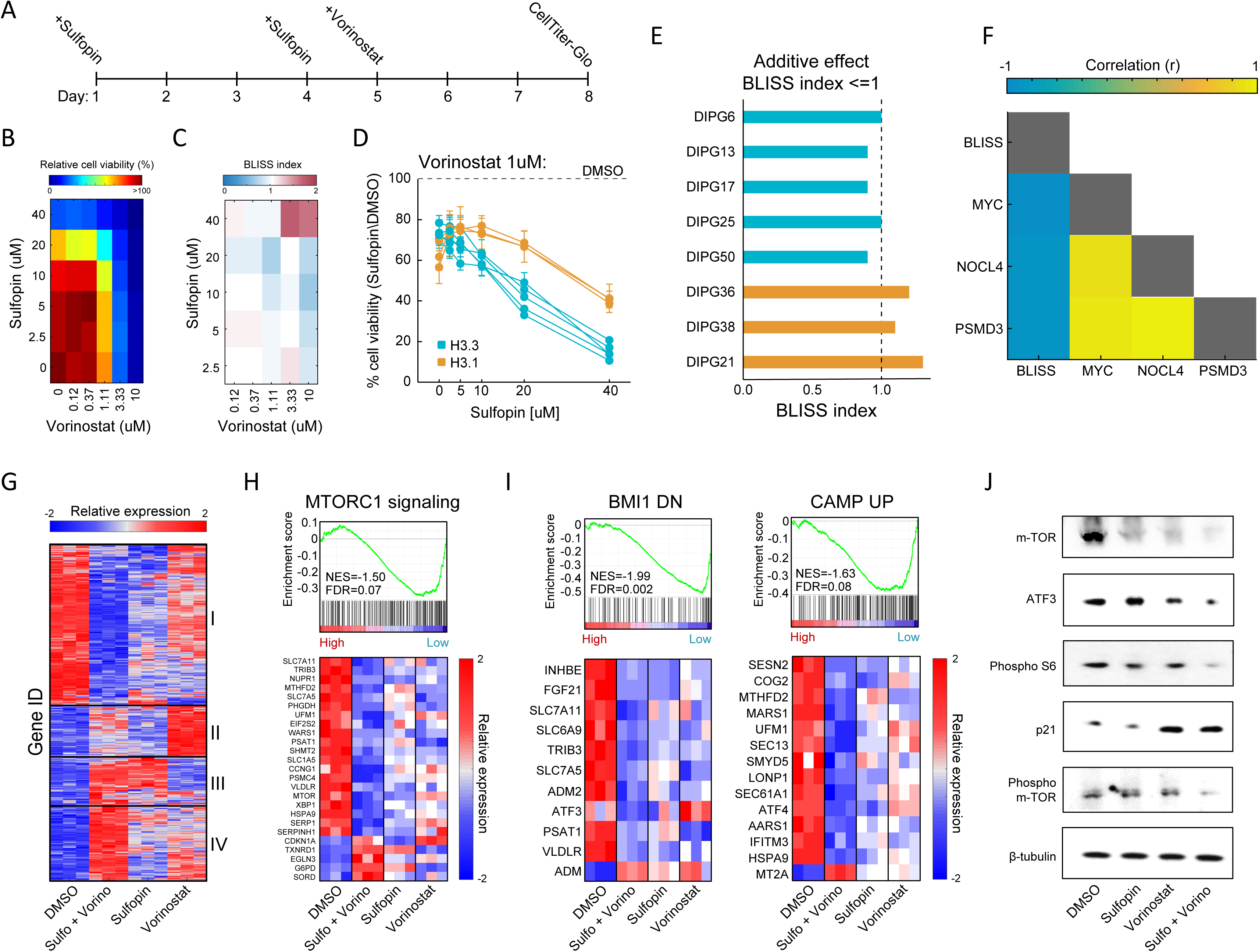
Combination of Sulfopin and Vorinostat elicits robust downregulation of oncogenic pathways. (A) Timeline demonstrating the treatment protocol for the combination of Sulfopin and Vorinostat. (B) Percentage of cell viability, as measured by CellTiterGlo of SU-DIPG13 cells treated with Sulfopin and Vorinostat at the indicated concentrations, compared to DMSO. (C) BLISS index measured as the ratio between the observed and the expected effect of the combination of Sulfopin and Vorinostat, for each pair of concentrations, in SU-DIPG13. Synergy: Bliss <1 ,Additive: Bliss=1, Antagonist: Bliss>1. (D) Cell viability as measured by CellTiterGlo, of eight DMG cultures treated with Sulfopin (0uM, 2.5uM, 5uM, 10uM, 20uM and 40uM) and Vorinostat (1uM), compared to DMSO. H3.3-K27M and H3.1-K27M cultures are indicated in blue and orange, respectively. Mean±SD of two technical replicates is shown. H3.3-K27M cells showed higher sensitivity to the combined treatment compared to H3.1-K27M cells. (E) The BLISS index of the combination of Sulfopin (10uM) and Vorinostat (1uM), in the indicated cultures. An additive effect was detected in all the H3.3-K27M cultures at this set of concentrations. (F) Pearson correlation coefficient matrix of BLISS index of the combined treatment (Sulfopin (10uM) and Vorinostat (1uM)) and mRNA levels of MYC and its target genes, in the eight DMG cultures tested. mRNA levels were measured by RT-qPCR (Fig. S2F-G). Blue and yellow colors indicate negative or positive correlation, respectively. Negative correlation was detected between the BLISS indexes and the expression levels of MYC and its target genes. (G) Unsupervised hierarchical clustering of expression levels of 620 significantly DE genes detected in SU-DIPG13 cells treated with either Sulfopin (10uM, 8 days), Vorinostat (1uM, 72 hours), the combination of Sulfopin and Vorinostat or DMSO. Gene expression rld values (log2 transformed and normalized) were standardized for each gene (row) across all samples. Color intensity corresponds to the standardized expression, low (blue) to high (red). Clusters 1 and 4 demonstrate additive transcriptional patterns associated with the combined treatment. (H) Top: Gene Set Enrichment Analysis (GSEA) on SU-DIPG13 treated with combination of 10uM Sulfopin and 1uM Vorinostat compared to DMSO, showing significant downregulation of mTORC1 signaling (‘HALLMARK_MTORC1_SIGNALING’ geneset^119^) in the combined treatment. NES: Normalized Enrichment Score. FDR: false discovery rate. Bottom: Expression levels of significantly DE genes detected in the combined treatment compared to DMSO that are part of the mTORC1 signaling geneset. *MTOR* gene was added manually to the heat-map. Heatmaps were generated as described in B. (I) Top: Gene Set Enrichment Analysis (GSEA) on SU-DIPG13 treated as in C, showing significant downregulation of the epigenetic BMI-1 pathway and the oncogenic cAMP pathway in the combined treatment (BMI1_DN.V1_UP; CAMP_UP.V1_UP; MSigDB C6 oncogenic signature^120,121^). Bottom: Expression levels of significantly DE genes detected in the combined treatment compared to DMSO that are part of the BMI-1 and cAMP genesets. Heatmaps were generated as in described B. (J) Western blot of SU-DIPG13 treated either with Sulfopin (10uM, 8 days), Vorinostat (1uM, 72 hours), the combination of Sulfopin and Vorinostat or DMSO, using the indicated antibodies. β-tubulin is used as loading control.

To gain more insights on the molecular mechanisms underlying this effect on viability, we profiled the transcriptome of SU-DIPG13 cells (H3.3-K27M) treated with each of the drugs alone and the combined treatment (Fig. S2J). A total of 494 genes were significantly differentially expressed (DE) in cells treated with the drug combination, compared to only 145 and 167 in cells treated solely with Sulfopin or Vorinostat, respectively (Fig. S2K-L). Hierarchical clustering of all 620 DE genes confirmed strong additive effect of the combined treatment on gene expression, primarily demonstrated by clusters 1 and 4 (Fig. 2G). Cluster 1, consisting of approximately half of the DE genes, was strongly downregulated by the combined treatment, compared to single-agent treated cells. This cluster included genes that play crucial roles in glioma progression such as *MTOR*, *ATF3*, *CREB3 and TGFA*^49–52^ (Fig. S2M). Moreover, the mTORC1 signaling pathway was highly enriched within the genes comprising cluster 1 (Fig. S2N). Importantly, GSEA analysis revealed that this pathway was significantly enriched in downregulated genes following the combined treatment (Fig. 2H and S2O). Of note, additional epigenetic and oncogenic pathways such as BMI1, cAMP and IL-6/JAK/STAT3 were significantly downregulated in cells treated with the drug combination^53–55^ (Fig. 2I and S2O). Reduction in mTOR signaling (mTOR, phospho-mTOR and phospho-S6^56^) and ATF3 was further validated by western blot and RT-qPCR analysis (Fig. 2J, S2P). Importantly, we also confirmed that the combined treatment with Sulfopin and Vorinostat led to induction of p21 (CDKN1A), a prominent tumor suppressor promoting cell cycle arrest and regulated by mTOR signaling^57–59^ (Fig. 2H, J). Finally, exploring mTOR expression in all the tested DMG cultures, revealed a strong negative correlation with the BLISS indexes of the combined treatment (Fig. S2Q), suggesting mTOR as a potential biomarker of response for the combined treatment of Sulfopin and Vorinostat. Overall, our data revealed that the combined inhibition of HDACs and MYC results in robust downregulation of prominent oncogenic pathways.

### Sulfopin attenuates HDACi-mediated accumulation of H3K27ac on genomic regions associated with oncogenic pathways

We next examined epigenetic features that may underly the inhibition of oncogenic pathways by the combined treatment (keeping the same experimental set-up as shown in figure 2A). Inhibition of histone deacetylases is expected to increase and alter genome-wide histone acetylation patterns, including the H3K27ac mark associated with active promoters and enhancers^60^. Indeed, global quantification by single-molecule measurements of H3K27-acetylated nucleosomes, as well as western blot analysis, confirmed HDACi-mediated increase in H3K27ac global levels (Fig. 3A-C, S3A). Interestingly, treatment with Sulfopin alone had the opposite effect, reducing the percentage of acetylated nucleosomes (Fig. 3B-C, S3A). As acetylation is coupled with transcription activation, this is in line with the function of Sulfopin as an inhibitor of MYC transcriptional activity^26,61^. As a result, the global HDACi-mediated increase in histone acetylation was also attenuated in the combined treatment with Sulfopin (Fig. 3B-C, S3A).

**Figure 3:**
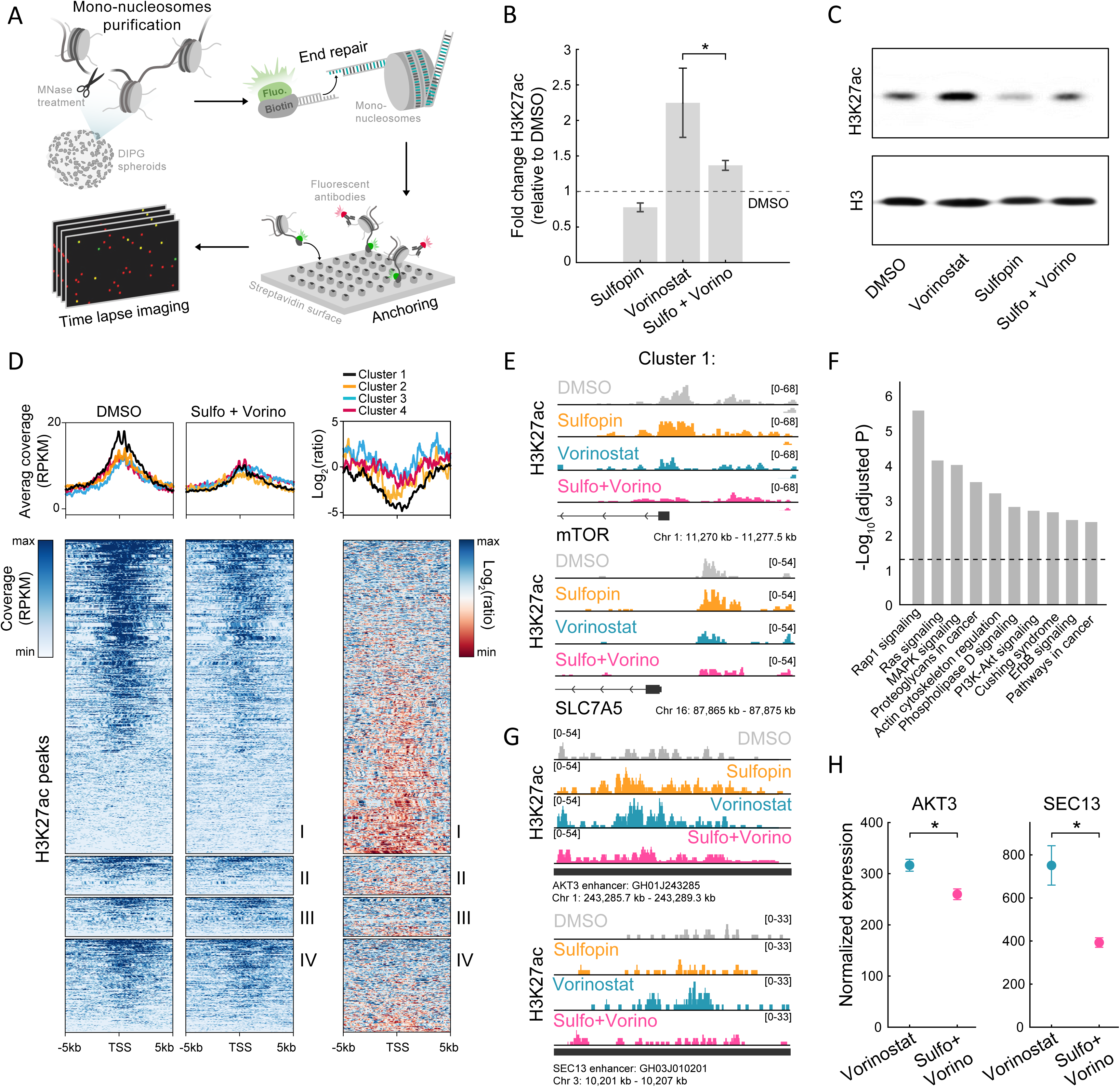
The combined treatment attenuates H3K27ac levels on oncogenic targets. (A) Scheme of the single-molecule imaging experimental setup^133^: cell-derived mono-nucleosomes are anchored in a spatially distributed manner on polyethylene glycol (PEG)-coated surface. Captured nucleosomes are incubated with fluorescently labeled antibodies directed against the H3K27ac modification. Total internal reflection fluorescence (TIRF) microscopy is utilized to record the position and modification state of each nucleosome. Time series images are taken to allow detection of maximal binding events. (B) Single-molecule imaging quantification of the percentage of H3K27ac nucleosomes, in SU-DIPG13 cells treated with either Sulfopin (10uM, 8 days), Vorinostat (1uM, 72 hours), or the combination of Sulfopin and Vorinostat, normalized to DMSO. Mean fold ± SE of at least two independent experiments is shown. H3K27ac global levels are lower in the combined treatment compared to cells treated solely with Vorinostat. *P < 0.05 (two sample t-test). (C) SU-DIPG13 cells were treated as in B, and analyzed by western blot using the indicated antibodies. (D) Left panel: Heatmap shows H3K27ac read coverage around the TSS (+/-5Kb) of the significantly DE genes shown in figure 2B, in SU-DIPG13 cells treated with the combination of 10uM Sulfopin and 1uM Vorinostat versus DMSO. Average coverage is shown on top. Color intensity corresponds to the standardized expression. Clusters 1-4 are indicated. Right panel: The log2 ratio of H3K27ac read coverage in SU-DIPG13 cells treated with the combination of 10uM Sulfopin and 1uM Vorinostat vs. DMSO was calculated. Heatmap shows the ratio around the TSS (+/-5Kb) of the significantly DE genes shown in figure 2B, and average coverage is shown on top. Color intensity corresponds to the ratio between samples, low (red) to high (blue). Clusters 1-4 are indicated, with cluster 1 presenting the strongest local decrease in H3K27ac following the combined treatment compared to DMSO. (E) IGV tracks of *MTOR* and *SLC7A5* gene promoters, showing H3K27ac coverage in SU-DIPG13 cells treated as indicated. (F) Functional enrichment analysis of the genes linked to enhancers (top targets of high confident enhancers) marked with H3K27ac exclusively in SU-DIPG13 cells treated with Vorinostat, and not in the combined treatment. gProfiler algorithm^126^ was used to calculate enrichment against the KEGG pathways DB^128^. Dashed line denotes adjusted p-value = 0.05. Genes associated with Vorinostat-unique enhancers are enriched for oncogenic signaling pathways. (G) IGV track of *AKT3* and *SEC13* linked enhancers, showing H3K27ac coverage in SU-DIPG13 cells treated with 1uM Vorinostat or the combination of 10uM Sulfopin and 1uM Vorinostat. (H) Normalized expression levels of *AKT3* and *SEC13* genes in SU-DIPG13 cells treated as in G. Mean ± SD of three technical repeats is shown. *P < 0.05 (two-sample t-test).

To profile changes in the genomic distribution of H3K27ac and H3K27me3, we applied Cut&Run in SU-DIPG13 cells treated with Vorinostat alone or in combination with Sulfopin. We first examined the promoters of genes differentially expressed in the combined treatment (as clustered in Fig. 2G). Genes comprising cluster 1 showed higher basal expression levels compared to genes from cluster 4, and these expression differences were also reflected in higher H3K27ac at these genes’ promoters (Fig. S3B-C). Moreover, in accordance with the reduction in RNA expression levels of genes comprising cluster 1, the combined treatment led to the strongest local reduction of H3K27ac levels associated with their promoters (Fig. 3D). Specifically, we found that several of the oncogenes that comprise cluster 1, such as mTOR, and mTOR regulator-SLC7A5, RELB and COPB2, showed decreased levels of H3K27ac at their promoters following the combined treatment^62–64^ (Fig. 3E, S3D). Cluster 1 genes also gained H3K27me3 on their promoters upon the combined treatment, supporting their epigenetic repression following Sulfopin and Vorinostat treatment (Fig. S3E).

We further identified genomic regions in which histone acetylation is lost in a Sulfopin-dependent manner (i.e., H3K27ac peaks that are present in Vorinostat-treated cells, but are lost in the combined treatment with Sulfopin). Genome distribution analysis of these unique peaks revealed that the majority of them localize to distal regions (Fig. S3F) and more than 60% of them localize to annotated enhancers. Importantly, genes associated with these enhancers were strongly enriched for oncogenic pathways such as RAS, MAPK and PI3K-AKT signaling pathways^66–68^ (Fig. 3F, S3G). Specifically, the combined treatment reduced H3K27ac levels on several enhancers linked to genes involved in mTOR pathway and gliomagenesis, such as: *AKT3*, *JAK1* and *NFKB2* genes, that were shown to have critical role in malignant gliomas^69–73^, *SEC13* gene that indirectly activates mTOR^74,75^, and *CD93*, a key regulator in glioma^76^ (Fig 3G and S3H). Concomitant with the lower H3K27ac levels associated with these enhancers, we also observed lower expression of the associated genes (Fig. 3H and S3I). Taken together, the results show attenuation of HDACi-mediated accumulation of H3K27ac levels upon combined treatment with Sulfopin, associated with promoters and enhancers of prominent oncogenes. This is in-line with reduced expression of these oncogenic pathways, strengthening the therapeutic potential of the combined treatment.

### The combined treatment reduces tumor growth in a DMG xenograft mouse model

Finally, we examined whether the combined inhibition of HDACs and PIN1 elicits an additive effect also *in-vivo*. Thus, we established a H3-K27M DMG orthotopic xenograft model by injections of SU-DIPG13P* cells, engineered to express firefly luciferase, to the pons of immunodeficient mice^17^. Mice were given ten days to develop tumors, and then randomly assigned to four groups: control mice treated with DMSO, mice treated with Sulfopin or Vorinostat as single agents, and mice treated with the combination of Sulfopin and Vorinostat. Both drugs were given daily and tumor growth was monitored by in *in-vivo* bioluminescence imaging. In-line with our *in-vitro* results, we found significant reduction in tumor growth following the combined treatment compared to control DMSO treated mice (Fig. 4A-B). Importantly, the reduction was also seen when comparing to mice treated solely with Vorinostat, supporting the additive value of Sulfopin to the well-known anti-tumorigenic effect of HDACi in H3-K27M gliomas^17^ (Fig. 4A-B). Of note, mice from all groups showed signs of severe dehydration and general deterioration after 18 days of treatment, and thus we could not measure a survival benefit for the combined treatment. This rapid deterioration is likely a result of the aggressiveness of the transplanted tumors and does not represent side effects of the treatment, as mice from all groups, including the non-treated mice, showed similar signs of deterioration.

**Figure 4:**
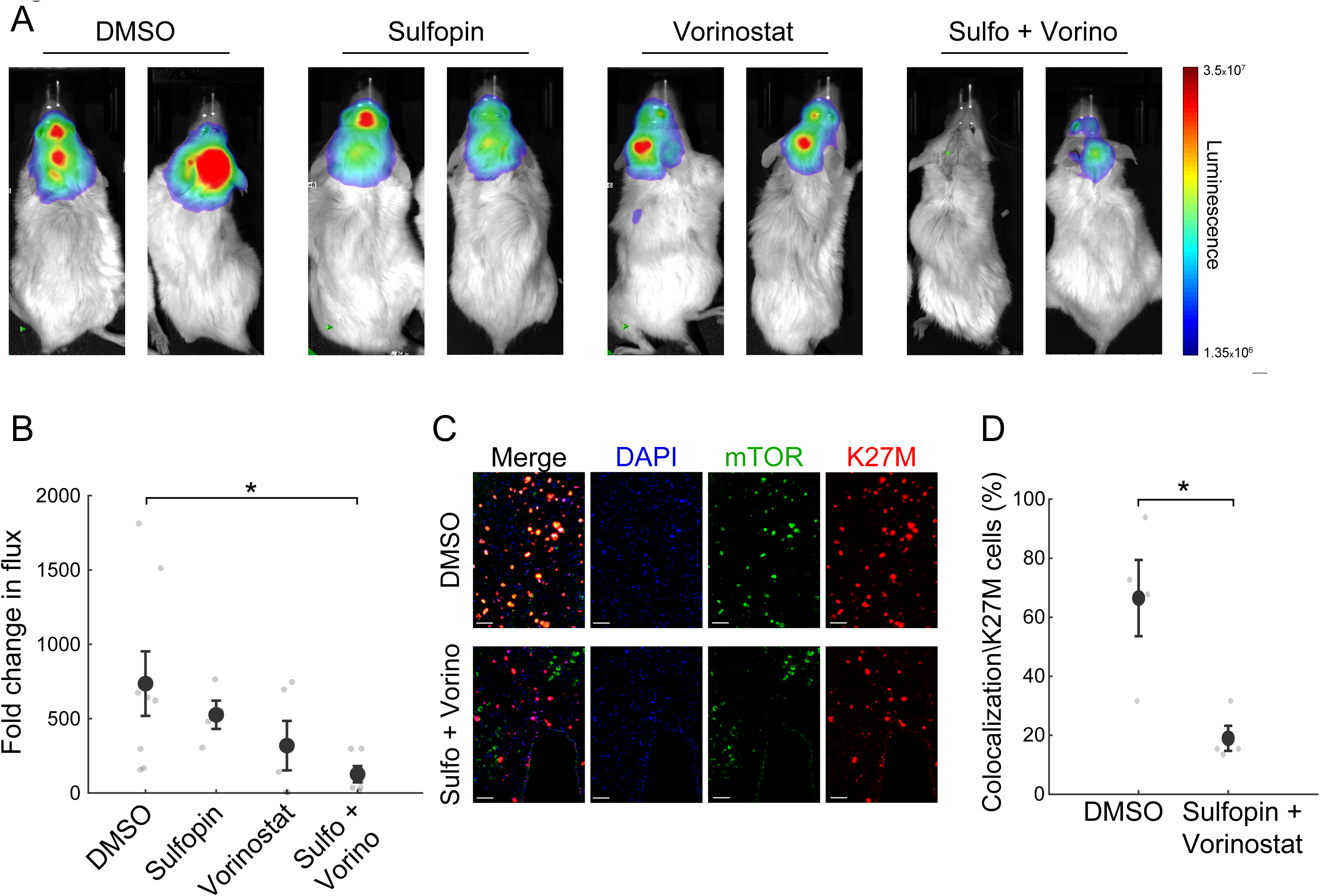
The combined treatment with Sulfopin and Vorinostat reduces tumor growth *in-vivo*. (A-B) SU-DIPG13P* cells were injected to the pons of immunodeficient mice to form tumors. Ten days post injection, mice were treated for 18 days with either DMSO, Sulfopin, Vorinostat, or the combination of Sulfopin and Vorinostat. (A) *In-vivo* bioluminescent imaging of DMG xenografts following 18 days of treatment. The heat map superimposed over the mouse head represents the degree of photon emission by DMG cells expressing firefly luciferase. (B) DMG xenograft tumor growth as measured by change in bioluminescent photon emission following 15 days of treatment with either DMSO (n=8), Sulfopin (n=4), Vorinostat (n=5) or the combination of Sulfopin and Vorinostat (n=6). Data points represent the fold-change in maximum photon flux between day 3 and day 18 under treatment for each mouse. *P < 0.05 (two-tailed Mann-Whitney U-test). (C-D) Immunofluorescent staining of brain sections from mice treated with DMSO (n=4) or the combination of Sulfopin and Vorinostat (n=4). (C) Representative fluorescence images of H3-K27M (red) and mTOR (green). (D) Percentage of mTOR positive cells out of the total H3-K27M-positive cells (n=13-198 cells per FOV). H3-K27M positive cells show lower levels of mTOR following the combined treatment compared to DMSO. *P < 0.05 (two-tailed t-test).

To link our *in-vitro* transcriptomic analysis to the *in-vivo* effect on tumor growth, we assessed the percentage of H3-K27M cells that are positive to mTOR signal (double-positive), out of the total number of H3-K27M cells, by immunofluorescence staining. The combined treatment significantly reduced the fraction of the double-positive cells, compared to DMSO (Fig. 4C-D). Of note, mTOR signal in the mouse cells surrounding the tumor showed no significant differences between the DMSO and the combination-treated mice (Fig S4A-B). Overall, these results verify the potency of this treatment in restricting oncogenic pathways also *in-vivo*, and the potential clinical benefit of PIN1 and HDAC combined inhibition in H3-K27M-mutant DMGs.

## Discussion

Despite decades of efforts to improve treatment for children diagnosed with H3-K27M-mutant DMGs, the standard of care remains solely radiation, with no significant improvement in overall survival^1,2,79^. High throughput drug screens in patient-derived cells identified HDAC inhibition as a promising therapeutic strategy, and ongoing clinical trials revealed preliminary evidence of clinical benefit^38,80^ (NCT02717455, NCT03566199). However, emerging resistance to HDACi in glioma cells^17^, as well as limited therapeutic benefits of HDACi treatment as a sole agent^81,82^, highlight the necessity to discover additional therapy modalities that could prolong survival and relieve symptoms.

In H3-K27M-mutant DMGs, as well as in many other malignancies, high MYC activity is correlated with poor prognosis and therefore considered as an attractive therapeutic target^21,83^. Nevertheless, direct targeting of MYC is highly challenging, functional and structural wise. MYC signaling, while amplified in cancer cells, is also essential to normal cell function^84^. Moreover, as a transcription factor, MYC lacks traditionally druggable binding pockets, hindering the development of direct small molecule binders^25^. To overcome this, in this study we exploited Sulfopin, a novel, potent and selective PIN1 inhibitor^26^. Pre-clinical studies have shown *in-vivo* activity and negligible toxicity of Sulfopin in murine models of PDAC and neuroblastoma^26^. While PIN1 is not overexpressed in DMG tumors, we show that Sulfopin, although not inhibiting MYC directly, downregulates MYC target genes in patient-derived DMG cells and affects cell survival in a H3-K27M-dependant manner. These results are in line with the contribution of MYC signaling to H3-K27M-driven tumorigenesis^23,24^, underscoring the therapeutic potential of Sulfopin for these aggressive brain tumors. Notably, despite a significant reduction in tumor size in-vivo, the combined treatment did not increase mice survival. This is perhaps due to the relatively large tumors already formed at the onset of treatment, leading to rapid deterioration of mice in all experimental groups. Thus, further optimization of the modeling system and therapeutic regime is needed.

We showed that combining Sulfopin treatment with the HDAC inhibitor Vorinostat reduced cell viability by 80% in several tumor-derived DMG lines originating from different patients. Importantly, the combination resulted in an additive effect not only on cell viability, but also on transcriptional programs. A key oncogenic pathway that was robustly downregulated in the combined treatment, *in-vitro* and *in-vivo,* is the mTOR signaling pathway. Preclinical and clinical data support a role for mTOR in gliomagenesis^85^. Specifically, AKT gain or PTEN loss is detected in approximately 70% of DMG tumors, and a recent study showed high expression of mTOR in DMGs. Thus, the PTEN/AKT/mTOR pathway is suggested to play a key role in this disease^86–89^. Attempts to restrict mTOR signaling using first generation mTOR inhibitors primarily inhibited mTORC1, which often caused upregulation of mTORC2^90^. The combined treatment with Sulfopin and Vorinostat led to robust inhibition of several target proteins in the mTOR signaling pathway, likely via alternative pathways. One potential candidate is the mTOR-dependent cell-growth through MYC regulation, which may be impaired by Sulfopin treatment, inhibiting MYC transcriptional activity^91–94^. Of note, drug agents targeting both mTORC1 and mTORC2 were shown to be highly effective in DMG cells as well as in murine models^95,96^.

Expression of the H3-K27M oncohistone is associated with a prominent increase in H3K27ac levels^11–13,16^. Among the loci that accumulate this modification are repetitive elements and endogenous retroviruses (ERVs), resulting in their aberrant expression, potentially inducing innate immune responses^12^. However, H3-K27M cells do not show an increase in interferon stimulated genes (ISGs), preassembly due to high levels of MYC, which is potent antagonist of interferon responses^12,97^. Thus, dual targeting of both HDAC and MYC pathways has the potential of inducing robust expression of ISGs, thus promoting tumor cell death mediated by viral-mimicry^98^.

The high levels of H3K27ac in H3-K27M cells was also linked to activation of super-enhancers, which regulate the expression of oncogenes and genes that are associated with an undifferentiated state^79^. Here, we show that H3-K27M cells are vulnerable to transcriptional disruption mediated by the combined treatment with Sulfopin and Vorinostat. While treating cells with Vorinostat alone increased H3K27ac levels, as expected, the combined treatment attenuated the accumulation of this mark on promoters and enhancers of prominent oncogenes, resulting in their decreased expression. This Sulfopin-mediated attenuation could potentially result from downregulation of MYC targets, as it was recently shown that MYC overexpression results in H3K27ac gain on super-enhancers^99^.

In summary, we propose a novel therapeutic strategy targeting epigenetic and transcriptional pathways in H3-K27M-mutant DMG tumors. HDAC inhibition, in combination with the novel MYC inhibitor, inhibited oncogenic transcriptional signatures and reduced tumor growth *in-vivo*. Our findings emphasize epigenetic dependencies of H3-K27M driven gliomas and highlight the therapeutic potential of combined treatments for this aggressive malignancy.

## Acknowledgments

We thank H. Keren-Shaul, D. Robbins, Y. Elazari and N. Adler (G-INCPM, WIS) for their help with NGS; N. Morris for her help with conducting part of the cell viability assays; C. Raanan, and M. Zerbib for their contribution in establishing the DIPG mouse model; I. Savchenko for his help with immunohistochemistry; R. Gabizon for generously providing Sulfopin for *in-vitro* and *in-vivo* experiments and helping with Sulfopin handling; S. Fishilevich for providing the GeneHancers data; L. Segev for computational framework for single-molecule image analysis; N. Harpaz and O. Beresh for helping with cultures maintenance; O. Griess for helping with transcriptomic protocols; S. Gillespie for providing valid information on DMG cells growth and maintenance; and I. Ulitsky for providing the pAG-MNase enzyme. We are thankful to O. Golani for helping with the immunohistochemistry image analysis. We are grateful to M. Monje for generously sharing with us the SU-DIPG13, SU-DIPG13P*, SU-DIPG6, SU-DIPG17, SU-DIPG25, SU-DIPG36, SU-DIPG38, SU-DIPG21 and SU-DIPG50 cells. We are grateful to N. Jabado for her kind gift of the isogenic BT245 and SU-DIPG13 cultures. We would like to express our sincere gratitude to D. Deitch for producing part of the graphs, designing the figures and illustrations, and helping with statistical analysis. **Funding:** E.S. is an incumbent of the Lisa and Jeffrey Aronin Family Career Development chair and is also supported by Henry Chanoch Krenter Institute for Biomedical Imaging and Genomics. This research was supported by grants from the European Research Council (ERC801655), The Israel Science Foundation (1881/19), Emerson Collective, and The Israel Cancer Research Fund: Research Career Development Award.

## Author contributions

D.A, N.F and E.S. designed the study and wrote the manuscript; D.A conducted the experiments; R.O and A.H established the xenograft mice model and performed *in-vivo* experiments; B.D, D.A, N.F and E.S performed bioinformatics analysis. L.F.A, R.O, G.R and D.A conducted the *in-vivo* image analysis; H.B and A.P conducted part of the viability assays; N.L suggested and provided Sulfopin and valuable usage information.

## Declaration of interests

N.L. is an inventor on a patent describing Sulfopin (US 2021/0332024 A1).

## Methods

### Data availability

All sequencing data is deposited in NCBI’s Gene Expression Omnibus (GEO) and available through GEO series accession number GSE221614.

### Cell cultures

All cell lines were maintained at 37°C with 5% CO2. Exclusion of Mycoplasma contamination was monitored and conducted by test with EZ-PCR kit (Biological Industries # 20-700-20).

### Glioma cultures

DIPG derived cells, SU-DIPG13 (H3.3-K27M, female), SU-DIPG6 (H3.3-K27M, female), SU-DIPG17 (H3.3-K27M, male), SU-DIPG25 (H3.3-K27M, female), SU-DIPG50 (H3.3-K27M, sex unknown), SU-DIPG36 (H3.1-K27M, female), SU-DIPG38 (H3.1-K27M, female), SU-DIPG21 (H3.1-K27M, male) and SU-DIPG13P* were generated in the lab of Dr. Michelle Monje, Stanford University^17^. SU-DIPG13P* are a subclone of the patient-derived SU-DIPG13 pons culture that demonstrates more aggressive growth in vivo^38,79^.

K27M-KO SU-DIPG13, BT245-KO (sex information is unavailable) and corresponding control clones were generated in the lab of Prof. Nada Jabado, McGill University, as previously described by Krug et al^12^.

Cells were cultured in Tumor Stem Media (TSM) consisting of a 1:1 mixture of DMEM/F12 (Invitrogen, 31330038) and Neurobasal -A (Invitrogen, 10888022) media, with 1:100 addition of HEPES Buffer Solution (Invitrogen 15630-080), MEM Sodium Pyruvate Solution (Invitrogen, 11360-070), MEM Non-Essential Amino Acids Solution (Invitrogen, 11140-050), GlutaMAX-I (Invitrogen, 35050-061) and Antibiotic-Antimycotic (Invitrogen,15240-096). The following additive were added freshly: B27 -A (1:50, Invitrogen, 12587010), human FGF (20 ng/ml, Shenandoah Biotechnology, 100-146), human EGF (20 ng/ml, Shenandoah Biotechnology, 100-26), human PDGF-AA (20 ng/ml, Shenandoah Biotechnology, 100-16), human PDGF-BB (20 ng/ml, 100-18, Shenandoah Biotechnology) and heparin (10 ng/ml, Stemcell Technologies, 07980). Doubling times were computed using: http://www.doubling-time.com/compute.php.

### Astrocytic-like cells culture

CRL-1718 cells, an astrocytic-like cells isolated from the brain of patient with astrocytoma were kindly provided by Dr. D. Michaelson, Tel-Aviv University, Israel. Cells were cultured in RPMI media (Biological Industries, # 01-101-1A) supplemented with 10% heat inactivated Fetal Bovine Serum (FBS), 1 mM L-glutamine and 1% penicillin/streptomycin solution (all from Biological Industries).

### Isolation of total RNA, reverse transcription and RT-qPCR

RNA was isolated from 2 million cells using the NucleoSpin kit (Macherey Nagel, #740955.50). 1ug of each RNA sample was reverse-transcribed using Moloney Murine Leukemia Virus reverse transcriptase (M-MLV-RT, Promega, #M1701) and random hexamer primers (Thermo Scientific, #SO142). Real-time qPCR was performed using KAPA SYBR FAST mix (Kapa Biosystems, #KK4660) with a StepOne real-time PCR instrument (Applied Biosystems). For each gene, standard curve was determined and the relative quantity was normalized to GAPDH mRNA (Table S2).

### Isolation of total RNA, bulk MARS-Seq library preparation and sequencing

RNA was isolated from SU-DIPG cells treated as indicated in Table S4a, using the NucleoSpin kit (Macherey Nagel, 740955). A bulk adaptation of the MARS-Seq protocol^100,101^ was used to generate RNA-Seq libraries for expression profiling. Briefly, 30 ng of input RNA from each sample was barcoded during reverse transcription and pooled. Following Agencourct Ampure XP beads cleanup (Beckman Coulter, A63880), the pooled samples underwent second strand synthesis and were linearly amplified by T7 in vitro transcription. The resulting RNA was fragmented and converted into a sequencing-ready library by tagging the samples with Illumina sequences during ligation, RT, and PCR. Libraries were quantified by Qubit (ThermoFisher Scientific) and TapeStation (Agilent) as well as by qPCR for GAPDH housekeeping gene as described below. Sequencing was done on a Nextseq 75 cycles high output kit (Illumina, 20024906).

### Bulk MARS-seq analysis

A median of 10.5 to 20 million reads were obtained for each dataset (Table S4a). MARS-seq analysis was preformed using the UTAP transcriptome analysis pipeline^113^. Reads were trimmed to remove adapters and low quality bases using cutadapt^105^ and mapped to the human genome (hg38, UCSC) using STAR v2.4.2a18^114^ (parameters: –alignEndsType EndToEnd, – outFilterMismatchNoverLmax 0.05, –twopassMode Basic, –alignSoftClipAtReferenceEnds No). In short, the pipeline quantifies the 3’ of annotated genes (The 3’ region contains 1,000 bases upstream of the 3’ end and 100 bases downstream). Counting was done using HTSeq-count in union mode^115^. Genes having a minimum 5 UMI-corrected reads in at least one sample, were considered. Normalization of the counts and differential expression analysis was done using DESeq2^116^, and is provided in Table S4b-c (parameters: betaPrior=True, cooksCutoff=FALSE, independentFiltering=FALSE). Significantly differentially expressed genes were defined having adjusted p-value (with Benjamini and Hochberg procedure) <= 0.05, |log2FoldChange| >= 1 and baseMean >= 5. Heatmaps of gene expression were calculated using the log-normalized expression values (rld), with row standardization (scaling the mean of a row to zero, with standard deviation of 1). Clustering of gene expression (rld values) was done using Euclidian average linkage method, and visualized using Partek Genomics Suite 7.0 software (Partek Inc. 2020. Partek^®^ Genomics Suite^®^, Version 7.0). Boxplot was used to visualize gene expression values (rld) of each differentially expressed gene from clusters 1 and 4 (in each of the three replicates).

### Expression analysis from published data-sets

Gene expression data and histone mutation status of pediatric high grade gliomas^22^ was downloaded from PedcBioPortal (https://pedcbioportal.kidsfirstdrc.org/). For matched tumor-normal samples, RPKM values were obtain from Berlow et al^35^ (S2_Table). For boxplots the log2 (RPKM+1) or z-score values were used (ggpubr R package). Differences between groups were assessed using two-sided ‘t_test’ (rstatix R package). For heatmaps, gene expression values were obtained for genes from MsigDB signatures ‘HALLMARK_MYC_TARGETS_V1’, with MYC and PIN1 genes added manually. Heatmaps of gene expression were calculated using RPKM values, with row standardization (scaling the mean of a row to zero, with standard deviation of 1).

### Functional enrichment analysis

Gene set enrichment analysis (GSEA) was performed as described in www.broadinstitute.org/software/gsea^117,118^. Genes detected by MARS-seq were pre-ranked according to their fold change (log2) in Sulfopin+Vorinostat vs. DMSO treated cells. Pre-ranked GSEA was applied using the Human MSigDB Collections H: hallmark gene sets^119^, and the C6: oncogenic signature gene sets^120–122^. Significantly downregulated gene-sets were selected using cut-off of FDR q-value ≤ 0.1.

Enrichr algorithm^123^ was used to compare significantly downregulated genes detected in the different treatments, against one or more of the following databases: the Molecular Signatures Database (MSigDB) hallmark^119^, ChEA^124^ or ENCODE TF ChIP^125^.

gProfiler algorithm^126^ was used to analyze genes that are putative targets of elite enhancers which overlap with Vorinostat-unique H3K27ac peaks (see Cut&Run analysis), against the KEGG^127,128^ or Wiki^129^ pathways databases.

### Cut&Run pulldown assay followed by high-throughput sequencing

Cut&Run assay was done as described in Skene and Henikoff, 2017^31,102^ with slight modifications as follows: Cells were harvested and counted, with 200,000 cells taken per reaction. Permeabilized cells, bound to Concanavalin A-coated beads (Bangs Laboratories, BP531), were mixed with individual primary antibody (Table S1) and incubated overnight at 4°C while rotating. Secondary antibody, anti-rabbit HRP, was used as a negative control. pAG-MNase enzyme (generated in the Department of Life Sciences Core Facilities, WIS, using Addgene plasmid 123461) was added to each sample followed by incubation step of 1 h at 4°C. Targeted digestion was done by 15 min incubation on ice block (0°C) under low salt conditions. DNA purification was done using Nucleospin gel and PCR clean-up kit (Machery-Nagel, 740609).

Libraries were prepared from 1-20ng of DNA as previously described in Blecher-Gonen et al^103^. Briefly, DNA Fragments were repaired by T4 DNA polymerase (NEB, M0203), and T4 polynucleotide kinase (T4 PNK, NEB, M0201) was used to add a phosphate group at the 5′ ends. An adenosine base was then added by Klenow fragments (NEB, M0212) to allow efficient ligation, using T4 quick ligase (NEB, M2200), of sequencing adapters, which contain a T-overhang. DNA fragments were amplified by PCR reaction (Pfu Ultra II fusion, Agilent Technologies, 600670), which also introduced the Illumina-P5 adapter at one end of the molecule. SPRI beads were used to purify proper sized DNA fragments after each enzymatic step. Libraries were quantified by Qubit (ThermoFisher Scientific) and proper fragment range (200-400bp) was verified by TapeStation (Agilent). Sequencing was done on a Next-Seq 500 instrument (Illumina, #20024904) using a V2 150 cycles mid output kit (paired end sequencing).

Paired-end reads of each sample (∼7.5M median reads per sample, Table S5) were preprocessed with cutadapt^105^, to remove adapters and low-quality bases (parameters: --times 2 -q 30 -m 20), following evaluation of quality with FastQC. Reads were mapped to human genome (hg38, UCSC) using Bowtie version 2.3.5.1^106^ (--local --very-sensitive-local --no-unal --no-mixed -- no-discordant --dovetail -I 10 -X 700). Nucleosome fragments at the length >120bp were selected from the remaining unique reads using picard-tools, and broad peaks were called using MACS2 against the HRP samples as background control (parameters: -f BAMPE --SPMR -- nomodel --extsize 100 --keep-dup auto -q 0.05). Analysis of genomic features was done using ChIPseeker^107^. Reads coverage on TSS, gene body regions, specific gene-signatures and CGI were visualized using ngs.plot^108^. Annotation of CGI were downloaded for UCSC table browser. Association of peaks with specific genes was done using GREAT^109^. Bigwig files were constructed from BAM alignments using deepTools2 suite^110^, by ‘bamCoverage’ command, using RPKM-normalization in 10bp bins. Heatmaps and profiles were constructed in ‘scale-regions’ mode around peak summits, with missingDataAsZero paramter using ‘plotHeatmap’ and ‘plotProfile’ commands. In the heatmap of H3K27ac ratio (Fig. 3D) coverage was calculated as the log2 between treatment and DMSO, on TSS of genes which were depicted as differentially expressed by MARS-Seq. RPKM-normalized coverage files were visualized on the genome using IGV (2.8.6). H3K27ac peaks were further distinguished into Vorinostat-unique peaks (i.e., H3K27ac peaks that are present in Vorinostat-treated cells, but are lost in the combined treatment with Sulfopin), and to shared peaks (i.e., H3K27ac peaks that are present both in Vorinostat and in Sulfopin+Vorinostat treated cells, with a minimum of 1bp overlap) using bedtools^111^. H3K27me3 peaks unique to Sulfopin treatment were defined similarly. To detect the overlap between the Vorinostat-unique peaks with enhancer regions, we used the GeneHancer database v5.9^112^. We required a 30% overlap of the peak length with a known enhancer categorized as “Elite” group, and excluded enhancers annotated to promoters regions.

### Cell viability assay by CelltiterGlo

For Sulfopin and Vorinostat combination experiments, 384-well plates were pre-plated with 2-fold dilution matrix of Vorinostat (MCE Cat#HY-10221) and Sulfopin (generous gift from Dr. Nir London’s Lab, Weizmann institute of science), in 5 concentrations ranging from 120 to 10,000 nM (Vorinostat) and 2500 to 40,000 nM (Sulfopin). 1400 DIPG cells/well were plated into each well containing 75ul of growth media, and subjected to drug treatment for 3 days (Vorinostat) or 8 days with pulse at day 4 (Sulfopin). Single agent titrations were run in parallel. Following the treatment, cells were subjected to cell viability assay using CellTiter-Glo assay (# G7572, Promega, USA) in accordance to the manufacture protocol. Luminescence signal was detected by luminescence module of PheraStar FS plate reader (BMG Labtech, Ortenberg, Germany). Viability data was analyzed using GeneData 15 (Switzerland).

For generating Sulfopin dosage curves in DIPG versus Astrocytes cells, the appropriated amounts of Sulfopin or DMSO (control) were transferred using Echo Liquid Handler 550 (Beckman, USA) into white/white 384-well TC plate (Greiner, #781080). 1400 cells/well were dispensed by MultiDrop 384 dispenser (Thermo Scientific, USA) into 384-well plates. After 4 days, cells were subjected to cell viability assay using CellTiter-Glo assay (# G7572, Promega, USA) in accordance to the manufacture protocol. Luminescence signal was detected by luminescence module of PheraStar FS plate reader (BMG Labtech, Ortenberg, Germany). Viability data was analyzed using GeneData 15 (Switzerland).

For all the other viability assays, cells were plated at a density of 2500 cells/well in 96-well plates and subjected to drug treatment for 3 days (Vorinostat-MCE Cat#HY-10221, GSK-J4-Selleck Cat#S7070, EPZ-6438-Selleck Cat#S7128) or 8 days (Sulfopin, generous gift from Dr. Nir London’s Lab, and MM-102-Selleck Cat#S7265) (Table S3). At least two replicates were measured for each treatment. When treated for 8 days drug was also added at day 4. Cell viability was measured by CelltiterGlo assay (G7571, Promega) according to the manufacturer’s instructions. Luminescence was measured by Cytation 5 plate reader and viability was compared to DMSO treated control cells.

At least two technical replicates were averaged in each experiment, and mean ± SE of all biological repeats was calculated. Two sample t-test on all technical replicates was used to determine significant difference between the treatments. Bonferroni correction was applied whenever more than two comparisons were made.

### IC50 quantification

Sulfopin IC50 values were determined by 4-parameter log-logistic dose response model using SynergyFinder 2.0^130^.

### BLISS index calculation

The Bliss index was calculated as the ratio between observed effect of combined drug treatment and expected effect (based on the calculated effect of combining single-drug treatment)^45,46^:

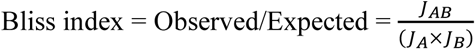

JA is the effect on growth of drug A only compared to DMSO

JB is the effect on growth of drug B only compared to DMSO

JAB is the effect on growth of drug A and B together compared to DMSO

Synergy: Bliss <1

Additive: Bliss=1

Antagonist: Blliss>1

### Western blot analysis

An equal number of cells from different treatments were suspended in Laemmli sample buffer (Bio-Rad, #1610747) containing 50mM DTT (Promega, #V3151), vortexed and heated up to 98°C. Samples were loaded onto Novex WedgeWell 4-20%, Tris-Glycine 4-20% gel (ThermoFisher Scientific, XP04205BOX) and electrophoresis was carried out in TG-SDS buffer (Bio-rad #1610732) for 1-1.5 hours at 120V. After electrophoresis, proteins were transferred to nitrocellulose membranes using transfer kit (Bio-rad, #1704156), followed by brief rinse in TBS (Bio-rad, #1706435) and blocking with 5% (w/v) milk powder in TBST (Tween20-Sigma, #P1379) for 60 minutes. The membranes were then rinsed in TBST and incubated with primary antibodies overnight at 4 °C with gentle rocking. Primary antibodies (Table S1) were diluted according to manufacturer instructions in TBST containing 5% (w/v) milk powder. The following day the membranes were rinsed in TBST, then incubated with HRP-conjugated secondary antibody (Table S1) diluted according to manufacturer instructions in a solution containing 5% (w/v) milk powder in TBST. The membranes were then rinsed in TBST and dipped in an ECL WB Detection Reagent (Bio-rad #1705061) prior to exposure. Membranes were imaged using a Bio-Rad ChemiDoc MP imaging system.

### Nucleosome preparation for single-molecule imaging

Nucleosomes for single-molecule imaging were extracted and labeled as described in Furth et al^16^. Briefly, 2–2.5 million cells were collected, washed once with PBS supplemented with protease inhibitors cocktail, 1:100, Sigma P8340), and HDAC inhibitors (20mM Sodium butyrate, Sigma 303410 and 0.1mM Vorinostat V-8477), and then resuspended in 0.05% IGEPAL (Sigma I8896) diluted in PBS (supplemented with inhibitors). Cell pellets were then lysed in parallel to chromatin digestion (100mM Tris-HCl pH 7.5, 300mM NaCl, 2% Triton® X-100, 0.2% sodium deoxycholate, 10mM CaCl2) supplemented with inhibitors and Micrococcal Nuclease (ThermoFisher Scientific, 88216). The suspension was incubated at 37°C for 10 min and the MNase reaction was inactivated by addition of EGTA at a final concentration of 20mM. Lysate was clear by centrifugation and nucleosomes were concentrated using an Amicon ultra-4 (Millipore, UFC810024). Inhibitors were supplemented following concentration. Nucleosomes labeling the following reaction was used: NEBuffer™ 2 (NEB B7202), inhibitors (as detailed above, 0.25 mM MnCl2, 33uM fluorescently labeled dATP (Jena Bioscience, NU-1611-Cy3), 33uM biotinylated dUTP (Jena Bioscience, NU-803-BIOX), 1.5ul of Klenow Fragment (3’→5′ exo-, NEB, M0212S) and 1.5ul of T4 Polynucleotide Kinase (NEB, M0201L). Samples were incubated at 37°C for 1h and then inactivated by addition of EDTA at a final concentration of 20mM. Nucleosomes were then purified on Performa Spin Columns (EdgeBio, 13266) followed by the addition of inhibitors.

### Surface preparation for single-molecule imaging

PEG-biotin microscope slides were prepared as described in Furth et al^16^. Ibidi glass coverslips (25mm×75mm, IBIDI, IBD-10812) were cleaned with (1) ddH2O (3X washes, 5 min sonication, 3X washes), (2) 2% Alconox (Sigma 242985, 20min sonication followed by 5X washes with ddH2O), (3) 100% Acetone (20min sonication followed by 3X washes with ddH2O). Slides were then incubated in 1M KOH solution for 30min while sonicated (Sigma 484016), followed by 3X washes with ddH2O. Slides were sonicated for 10 min in 100% HPLC EtOH (J.T baker 8462-25) and then incubated for 24 min in a mixture of 3% 3-Aminopropyltriethoxysilane (ACROS Organics, 430941000) and 5% acetic acid in HPLC EtOH, with 1 min sonication in the middle. Following washes with HPLC EtOH (3X) and ddH2O (3X) slides were dried and mPEG:biotin-PEG solution was applied (20mg Biotin-PEG (Laysan, Biotin-PEG-SVA-5000), 180mg mPEG (Laysan, MPEG-SVA-5000) dissolved in 1560ul 0.1M Sodium Bicarbonate (Sigma, S6297) on one surface followed by the assembly of another surface on top. Surfaces were incubated overnight in a dark humid environment and at the following day, were washed with ddH2O and dried. MS (PEG) 4 (ThermoFisher Scientific, TS-22341) was diluted in 0.1M of sodium bicarbonate to a final concentration of 11.7 mg/mL and applied on the surfaces for overnight incubation in dark humid environment. Surfaces were washed with ddH2O and dried.

### Single-molecule imaging of histone modifications

PEG-biotin coated coverslips were assembled into Ibidi flowcell (Sticky Slide VI hydrophobic, IBIDI, IBD-80608). Streptavidin (SIGMA, S4762) was added to a final concentration of 0.2mg/mL followed by an incubation of 10 min. TetraSpeck beads (used for image alignment, ThermoFisher Scientific, T7279) diluted in PBS were added and incubated on surface for at least 30 min. Labeled nucleosomes were incubated for 10 min in imaging buffer (10mM MES pH 6.5 (Boston Bioproducts Inc, NC9904354), 60mM KCL, 0.32mM EDTA, 3mM MgCl2, 10% glycerol, 0.1mg/mL BSA (Sigma, A7906), 0.02% Igepal (Sigma, I8896) to allow immobilization via biotin-streptavidin interactions and washed with imaging buffer. H3K27ac antibody was diluted in imaging buffer (1:2000) and incubated for 30 min. All positions (80– 100 fields-of-view (FOV) per experiment) were then imaged by a total internal reflection (TIRF) microscope by Nikon (Ti2 LU-N4 TIRF) every 10 min (10–15 cycles).

Image analysis was performed with Cell Profiler image analysis tools (http://www.cellprofiler.org/) as described in Furth et al^16^. Image analysis is done in two steps: (1) Time-lapse images of antibody binding events and TetraSpeck beads are aligned, stacked and summed to one image. Antibody spots and TetraSpeck beads spots were distinguished based on the size of the spot. (2) Stacked images are aligned to the initial images of the nucleosomes based on TetraSpeck beads location spots, and only binding events that align with nucleosomes are filtered and saved for further analysis. To evaluate random co-localization (negative control), each stacked image is aligned to a 90° flipped image of the initial nucleosomes. The use of TetraSpeck beads and a highly accurate microscope stage allowed alignment of images with shifts of up to 30 pixels. Nucleosomes are initially distributed on the surface in low density to minimize overlap between spots. Average percentage of modified nucleosomes over all fields of view (FOV) imaged was calculated in each experiment. Mean ± SE of three to five biological repeats was calculated. Two sample t-test on all biological repeats was used to determine significant difference between the treatments.

### Mouse injection and treatment

All animal experiments were conducted in accordance with approved institutional animal care and use committee protocols. Mice were housed and handled in a specific-pathogen-free, temperature-controlled (22°C ± 1°C) mouse facility on a reverse 12/12 h light/dark cycle, Animals were fed a regular chow diet et libitum. Single-cell suspension of SU-DIPG13P* or SU-DIPG13 cells were stereotactically injected into the pons of male NGS mice (NOD-SCID-IL2R gamma chain-deficient, The Jackson Laboratory) as previously described in Venkatesh et al^104^. Injection coordinates were: 0.8 mm posterior to lambda, 1 mm lateral to the sagittal suture, and 5 mm deep. Briefly, the skull of the mouse was exposed, and a small bore hole (0.5 mm) was made using a high-speed drill at the appropriate stereotactic coordinates. Approximately 300,000-400,000 cells in 3ul volume were injected at a speed of 0.3ul per minute into the pons with a 26-gauge Hamilton syringe; following 10 min pause, the needle was removed at a speed of 0.2 mm/minute. After closing the scalp, mice were placed on a heating pad and returned to their cages after full recovery. Mice were randomized to treatment groups, and treatment started 10 days (SU-DIPG13P*) or 5 weeks (SU-DIPG13) post-injection. Treatment was given daily for 18 days, by intraperitoneal injection according to the following groups: Vorinostat 200mg/kg daily (LC, V-8477-SU-DIPG13P*; MCE, HY-10221-SU-DIPG13), Sulfopin 40 mg/kg (gift from Dr. Nir London’s Lab), combination 200 mg/kg Vorinostst and 40 mg/kg Sulfopin, control mice received solvent 15% DMSO in 50% PEG400 (Sigma) in 0.15M NaCl.

### Tumor size imaging using IVIS

For bioluminescence imaging, mice received 100 μl D-luciferin (33 mg/ml; Perkin-elmer, Cat#122799) intraperitoneally 10 min before isoflurane anesthesia and were placed in LagoX Imaging System (Spectral Instruments Imaging).

The bioluminescence signal was measured using the ROI tool in LagoX software (Spectral Instruments Imaging). The fold change in total Photon flux (p/s) over time was used to compare tumor growth between the groups. Mann-Whitney U-test (Two-tailed) was used to determine significant difference between the DMSO and Sulfopin+Vorinostat treated mice.

### Mouse brain slides staining

Brains were collected, fixed in 4% paraformaldehyde, and embedded in paraffin blocks. For H3-K27M and mTOR staining, slides were de-paraffinized (1-Xylene –Bio-lab #242500; 2-Xylene; 3-Eth 100%-Bio-lab #05250501;4-Eth-96%; 5-Eth-70%;) and incubated in acetone (Bio-lab, #010352) at -20C. Slides were incubated in hydrogen peroxide blocking solution (200ml Methanol-Bio-lab #136805, 1% HCl-Bio-lab #084102, 3% H2O2-JT Baker #2186-01) for blocking endogenous peroxidase activity. Antigen retrieval was performed using citric acid buffer at pH-6, followed by blocking with 20% NHS and 0.5% triton in PBS. Rabbit anti mTOR (1:100, Table S1) was diluted in 2% NHS and 0.5% triton and was incubated overnight at room temperature. Slides were then incubated with HRP conjugated goat anti rabbit (1:100, Table S1) diluted 2% NHS for 1.5hr followed by OPAL 650 (1:500, Akoya FP1496001KT) reagent incubation for 15 min. Antibodies were stripped using microwave treatment with citric acid at pH-6, blocked and again incubated with rabbit anti H3-K27M (1:100 table S1) overnight at room temperature. Slides were then incubated with HRP conjugated goat anti rabbit (1:100, Table S1) diluted 2% NHS for 1.5hr followed by OPAL 570 (1:500, Akoya FP1488001KT) reagent incubation for 15 min. Slides were then washed with PBS and incubated with Hoechst (1:1000) for 1 minutes, followed by mounting with 25×75×1mm glass slides using Aqua-Poly/Mount Mounting Medium. All slides were scanned using the Panoramic MIDI slide scanner (3D Histech, Hungary) at x20 magnification, to receive a full scan of the tumor section.

For H3-K27M and mTOR co-localization analysis, double stained cells were counted using ImageJ. First, threshold based segmentation was used to detect positive cells in each of the channels and the image calculator feature was used to find the double positive cells. The total number of double-positive cells was then divided by to total number of H3-K27M-positive cells to calculate the percentage co-localization\H3-K27M cells. Similar-sized FOVs were used for all the images analyzed, and 13-198 H3-K27M-positive cells were detected per FOV (an average of 90 cells per FOV). For mTOR signal quantification in non-tumor cells (mice cells) a region of interest was selected in a location on the section with no positive H3-K27M cells. Threshold based segmentation was used to detect the percentage mTOR stained area.

### Statistical analysis

Unless noted otherwise, p values were determined using two-sample t-test and are indicated in the figure legends.

Pearson correlation coefficient was used to assess the relationship between all variables. Scatter plot was generated using ggscatter (R ggpubr package).

## Notes

### Summary of Updates

The main text was revised for clarification; Figure 1F has been revised; Panel S1D has been added.

