## Supplementary Materials for "Dual Targeting of Histone Deacetylases and MYC as Potential Treatment Strategy for H3-K27M Pediatric Gliomas"

Figures S1-S4

Table S1: Antibodies used throughout the study

Table S2: Oligonucleotides used throughout the study

Table S3: Drugs and inhibitors used throughout the study

Table S4a-c: Summary tables of MARS-seq libraries

Table S5: Summary table of Cut&Run libraries

Figure S1

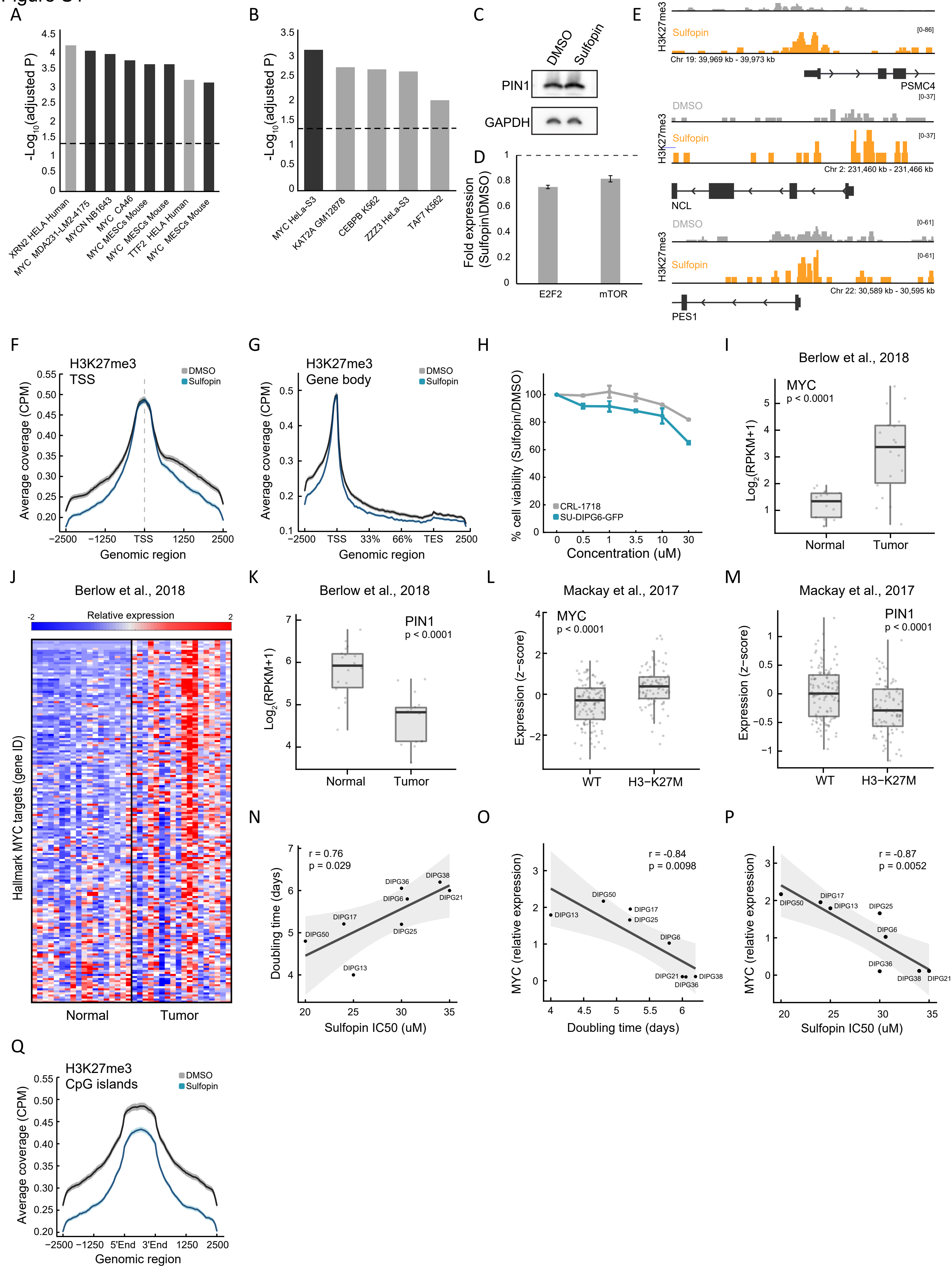
A
-Log10(adjusted P)
XRN2 HELA Human
MYC MDA231-LM2-4175
MYCN NB1643
MYC CA46
MYC MESCs Mouse
MYC MESCs Mouse
TTF2 HELA Human
MYC MESCs Mouse
B
MYC HeLa-S3
KAT2A GM12878
CEBPB K562
ZZZ3 HeLa-S3
TAF7 K562
C
DMSO
Sulfopin
PIN1
GAPDH
D
Fold expression (Sulfopin\DMSO)
E2F2
mTOR
E
H3K27me3
DMSO
Sulfopin
Chr 19: 39,969 kb - 39,973 kb
PSMC4
Chr 2: 231,460 kb - 231,466 kb
NCL
Chr 22: 30,589 kb - 30,595 kb
PES1
F
H3K27me3
TSS
Average coverage (CPM)
Genomic region
G
H3K27me3
Gene body
TES
H
% cell viability (Sulfopin/DMSO)
CRL-1718
SU-DIPG6-GFP
Concentration (uM)
I
Berlow et al., 2018
MYC
p < 0.0001
Log2(RPKM+1)
Normal
Tumor
J
Berlow et al., 2018
Relative expression
Hallmark MYC targets (gene ID)
K
Berlow et al., 2018
PIN1
p < 0.0001
L
Mackay et al., 2017
MYC
p < 0.0001
Expression (z-score)
WT
H3−K27M
M
Mackay et al., 2017
PIN1
p < 0.0001
N
r = 0.76
p = 0.029
Doubling time (days)
Sulfopin IC50 (uM)
O
r = -0.84
p = 0.0098
MYC (relative expression)
Doubling time (days)
P
r = -0.87
p = 0.0052
Q
H3K27me3
CpG islands
5'End
3'End

**Figure S1: The sensitivity of DMG cells to Sulfopin correlates with MYC expression levels**

(A-B) Functional enrichment analysis on significantly downregulated genes in SU-DIPG13 cells treated with 10uM Sulfopin for 12 hours, compared to DMSO. Enrichr[123] algorithm was used to compare the dataset against the ChEA[124] (A) and ENCODE TF ChIP[125] sets (B) databases. Dashed line denotes adjusted p-value = 0.05. Sulfopin downregulated genes are strongly enriched for MYC target genes as identified in different non-glioma cell lines (black bars).

(C) Western blot of SU-DIPG13 treated either with Sulfopin (10uM, 8 days) or DMSO, using the indicated antibodies.

(D) RT-qPCR analysis of E2F2 and mTOR genes in SU-DIPG6 cells treated with 10uM Sulfopin for 12 hours, compared to DMSO. Fold change between Sulfopin and DMSO treated cells was calculated and the mean ± SD of two technical repeats is shown.

(E) IGV track of the TSS region of three MYC targets, PSMC4, NCL and PES1, showing H3K27me3 coverage in SU-DIPG13 cells treated with 10uM Sulfopin for 8 days, compared to DMSO.

(F-G) H3K27me3 coverage over all TSS (F) and gene-body (G), in SU-DIPG13 cells treated as in C. Sulfopin does not affect H3K27me3 global deposition over these regions.

(H) Percentage of cell viability as measured by CellTiterGlo of SU-DIPG6-GFP and astrocytic-like cells (CRL-1718) treated with Sulfopin (0.5, 1, 3.5, 10, 30 uM) for 4 days, compared to DMSO. Mean ± SD of two technical repeats is shown. SU-DIPG6-GFP cells show higher sensitivity to Sulfopin compared to the astrocytes at all concentrations.

(I-K) Expression levels of MYC (I), 'MYC Targets V1' hallmark geneset[119] (J) and PIN1 (K) in 18 DMG tumors samples and their matched normal samples[35]. Box plots show center line as median, box limits as upper and lower quartiles, whiskers as minimum and maximum values. Individual data points are shown in grey circles. ***$P < 0.001$ (two-sample t-test). MYC and its target genes show higher expression in the DMG tumor samples, compared to their matched normal samples.

(L-M) Expression levels of MYC (K) and PIN1 (L) in WT and H3-K27M DMG tumors samples (n=201)[22]. Boxplot as in J. ***$P < 0.001$ (two-sample t-test).

(N) Doubling time (days) and Sulfopin IC50 levels measured in eight DMG cultures. Pearson correlation coefficient (r) is indicated. Grey area depicts 95% confidence interval for regression line. Positive correlation was detected between the two measures.

(O) Doubling time (days) and MYC expression levels measured in eight DMG cultures. Pearson correlation coefficient (r) is indicated. Grey area depicts 95% confidence interval for regression line. Negative correlation was detected between the two measures.

(P) MYC expression and Sulfopin IC50 levels measured in eight DMG cultures. Pearson correlation coefficient (r) is indicated. Grey area depicts 95% confidence interval for regression line. Negative correlation was detected between the two measures.

(Q) H3K27me3 coverage over CGIs, in SU-DIPG13 cells treated as in C. H3K27me3 signal is decreased in the Sulfopin-treated cells compared to the DMSO-treated cells.

### Figure S2

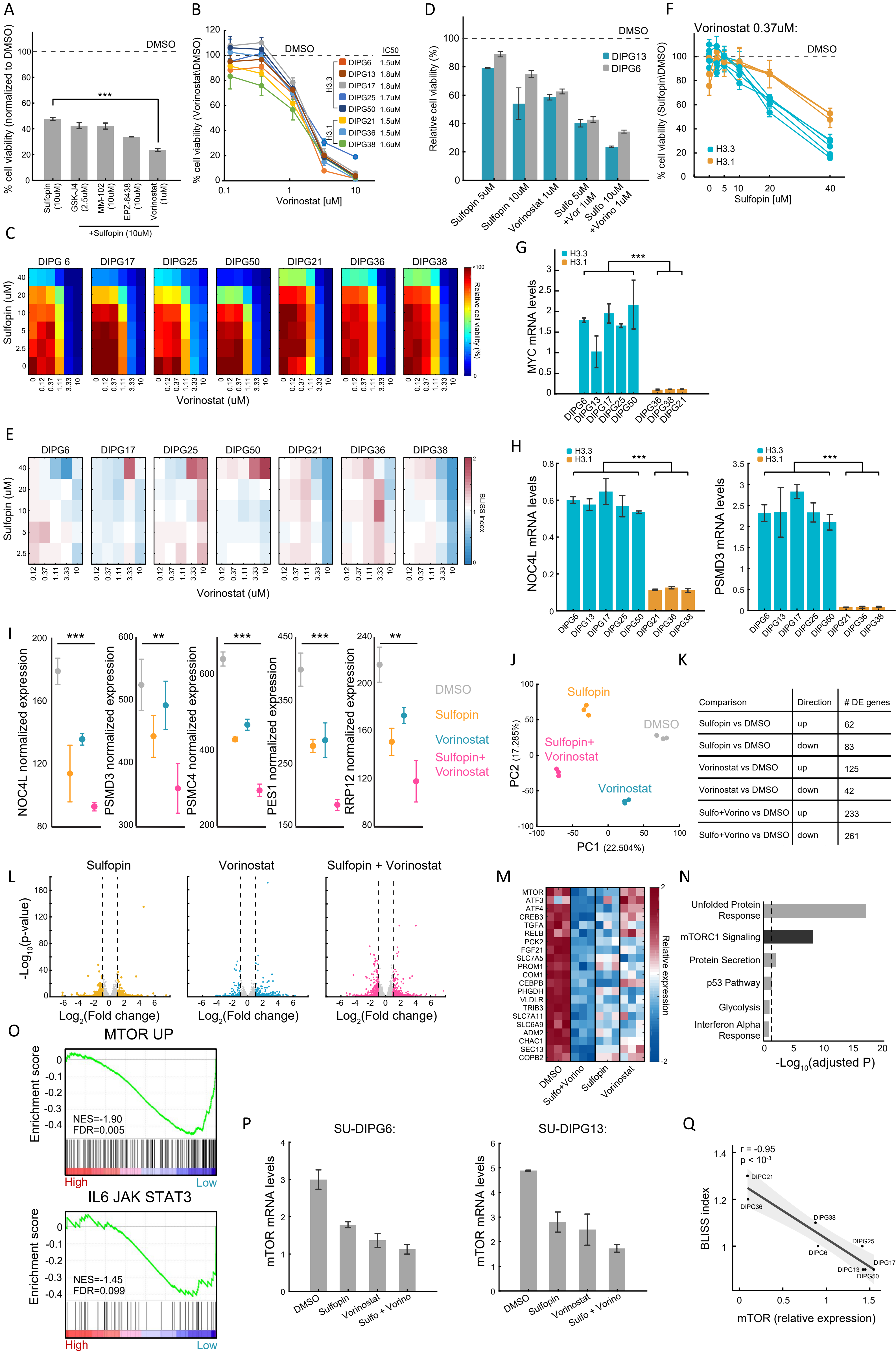
A
% cell viability (normalized to DMSO)
DMSO
***
Sulfopin (10uM)
GSK-J4 (2.5uM)
MM-102 (10uM)
EPZ-6438 (10uM)
Vorinostat (1uM)
+Sulfopin (10uM)
B
% cell viability (Vorinostat\DMSO)
DMSO
IC50
H3.3
DIPG6 1.5uM
DIPG13 1.8uM
DIPG17 1.8uM
DIPG25 1.7uM
DIPG50 1.6uM
H3.1
DIPG21 1.5uM
DIPG36 1.5uM
DIPG38 1.6uM
Vorinostat [uM]
D
Relative cell viability (%)
DMSO
DIPG13
DIPG6
Sulfopin 5uM
Sulfopin 10uM
Vorinostat 1uM
Sulfo 5uM +Vor 1uM
Sulfo 10uM +Vorino 1uM
F
Vorinostat 0.37uM:
% cell viability (Sulfopin\DMSO)
DMSO
H3.3
H3.1
Sulfopin [uM]
C
DIPG 6
DIPG17
DIPG25
DIPG50
DIPG21
DIPG36
DIPG38
Sulfopin (uM)
Vorinostat (uM)
Relative cell viability (%)
G
H3.3
H3.1
***
MYC mRNA levels
DIPG6
DIPG13
DIPG17
DIPG25
DIPG50
DIPG36
DIPG38
DIPG21
E
DIPG6
DIPG17
DIPG25
DIPG50
DIPG21
DIPG36
DIPG38
Sulfopin (uM)
Vorinostat (uM)
BLISS index
H
H3.3
H3.1
***
NOC4L mRNA levels
PSMD3 mRNA levels
I
NOC4L normalized expression
PSMD3 normalized expression
PSMC4 normalized expression
PES1 normalized expression
RRP12 normalized expression
DMSO
Sulfopin
Vorinostat
Sulfopin+ Vorinostat
J
Sulfopin
DMSO
Sulfopin+ Vorinostat
Vorinostat
PC2 (17.285%)
PC1 (22.504%)
K
Comparison | Direction | # DE genes
Sulfopin vs DMSO | up | 62
Sulfopin vs DMSO | down | 83
Vorinostat vs DMSO | up | 125
Vorinostat vs DMSO | down | 42
Sulfo+Vorino vs DMSO | up | 233
Sulfo+Vorino vs DMSO | down | 261
L
Sulfopin
Vorinostat
Sulfopin + Vorinostat
-Log10(p-value)
Log2(Fold change)
M
MTOR
ATF3
ATF4
CREB3
TGFA
RELB
PCK2
FGF21
SLC7A5
PROM1
COM1
CEBPB
PHGDH
VLDLR
TRIB3
SLC7A11
SLC6A9
ADM2
CHAC1
SEC13
COPB2
DMSO
Sulfo+Vorino
Sulfopin
Vorinostat
Relative expression
N
Unfolded Protein Response
mTORC1 Signaling
Protein Secretion
p53 Pathway
Glycolysis
Interferon Alpha Response
-Log10(adjusted P)
O
MTOR UP
Enrichment score
NES=-1.90
FDR=0.005
High
Low
IL6 JAK STAT3
NES=-1.45
FDR=0.099
P
SU-DIPG6:
SU-DIPG13:
mTOR mRNA levels
DMSO
Sulfopin
Vorinostat
Sulfo + Vorino
Q
r = -0.95
p < 10-3
BLISS index
mTOR (relative expression)
DIPG21
DIPG36
DIPG38
DIPG25
DIPG6
DIPG17
DIPG13
DIPG50

**Figure S2: The additive effect of the combination of Sulfopin and Vorinostat results in downregulation of oncogenic pathways**

(A) Percentage of cell viability, as measured by CellTiterGlo, of SU-DIPG13 treated with 10uM Sulfopin or with combined treatment of 10uM Sulfopin and the indicated agent (putative targets left-to right: JMJD3, MLL1, EZH2, HDAC), compared to DMSO. For Sulfopin and Vorinostat- mean ± SE of at least two independent experiments is shown. For all the other drugs-mean ± SD of three technical replicates is shown. In the combination of Sulfopin and Vorinostat, only ~20% of cells survive the treatment. ***P<0.001 (two-sample t-test).

(B) Cell viability, as measured by CellTiterGlo, of eight DMG cultures (SU-DIPG13, SU-DIPG6, SU-DIPG17, SU-DIPG25, SU-DIPG50, SU-DIPG36, SU-DIPG38 and SU-DIPG21), treated with the indicated concentrations of Vorinostat for three days, compared to DMSO. Mean±SD of two technical replicates is shown. Logarithmic scale is used for the x-axis.

(C) Percentage of cell viability, as measured by CellTiterGlo, of seven DMG cultures (SU-DIPG6, SU-DIPG17, SU-DIPG25, SU-DIPG50, SU-DIPG36, SU-DIPG38 and SU-DIPG21), treated with Sulfopin and Vorinostat at the indicated concentrations, compared to DMSO.

(D) Cell viability as measured by CellTiterGlo, of SU-DIPG13 and SU-DIPG6-GFP cells treated as indicated, relative to DMSO. Mean ± SE of two independent experiments is shown.

(E) BLISS index for each concentration of Sulfopin and Vorinostat, in the eight DMG cultures indicated in C. Synergy: Bliss <1 ,Additive: Bliss=1, Antagonist: Bliss>1. Additive effect was detected in the majority of the drug doses of the H3.3-K27M DMG cells, and in the higher dosages of Vorinostat in the H3.1-K27M cells.

(F) Cell viability as measured by CellTiterGlo, of eight DMG cultures treated with Sulfopin (0uM, 2.5uM, 5uM, 10uM, 20uM and 40uM) and Vorinostat (0.33uM), compared to DMSO. H3.3-K27M and H3.1-K27M cultures are indicated in blue and orange, respectively. Mean±SD of two technical replicates is shown. H3.3-K27M cells showed higher sensitivity to the combined treatment compared to H3.1-K27M cells.

(G-H) RT-qPCR analysis of MYC, and its target genes NOC4L and PSMD3, in eight DMG cultures. Mean ± SD of two technical repeats is shown. H3.3-K27M cells show higher expression compared to the H3.1-K27M cells. ***P < 0.001 (two-sample t-test).

(I) Normalized expression levels of the indicated MYC target genes in SU-DIPG13 cells treated as described in Fig. 2G. Mean ± SD of three technical repeats is shown. The expression of these genes is reduced following the combined treatment. **P < 0.01, ***P<0.001 (two-sample t-test).

(J) RNA sequencing was performed on SU-DIPG13 treated with either Sulfopin (10uM, 8 days), Vorinostat (1uM, 72 hours) or Sulfopin+Vorinosat (10uM, 8 days and 1uM, 72 hours), compared to DMSO. Principal component analysis (PCA) of all genes detected by RNA-seq is shown. Three technical repeats are shown.

(K) Table presenting the total numbers of significantly differentially expressed genes in each treatment compared to DMSO (adjusted p-value $\leq 0.05$, $|log2FoldChange| \geq 1$ and baseMean $\geq 5$).

(L) Volcano plots presenting all genes detected in each treatment compared to DMSO. Significantly differentially expressed genes are colored.

(M) Heat map presenting selected genes from cluster 1 (as shown in Fig. 2B) that were downregulated in the combined treatment compared to DMSO. Color intensity corresponds to the standardized expression, low (blue) to high (red). Only oncogenes and genes that are involved in glioma progression are shown.

(N) Functional enrichment analysis on genes comprising cluster 1 using Enrichr[123] algorithm comparing to the Molecular Signatures Database (MSigDB) hallmark geneset collection[119]. Dashed line denotes adjusted p-value = 0.05. mTORC1 signaling is highly enriched among cluster 1 genes.

(O) Gene Set Enrichment Analysis (GSEA) on SU-DIPG13 treated with combination of 10uM Sulfopin and 1uM Vorinostat compared to DMSO, showing significant downregulation of the mTOR oncogenic signature (MTOR_UP.N4.V1_UP; MSigDB C6 Oncogenic Signature[122]), and the IL-6/JAK/STAT3 signaling pathway (HALLMARK_IL6_JAK_STAT3_SIGNALING; MSigDB hallmark geneset collection[119]), in the combined treatment. NES: Normalized Enrichment Score. FDR: false discovery rate.

(P) RT-qPCR analysis of mTOR gene in SU-DIPG6 cells (left) and SU-DIPG13 (right), treated as indicated in Fig. 2G. Mean ± SD of two technical repeats is shown. The expression of mTOR is reduces following the combined treatment.

(Q) mTOR expression and BLISS index measured in 8 DMG cultures. Pearson correlation coefficient (r) is indicated. Grey area depicts 95% confidence interval for regression line. Negative correlation was detected between the two measures.

Figure S3

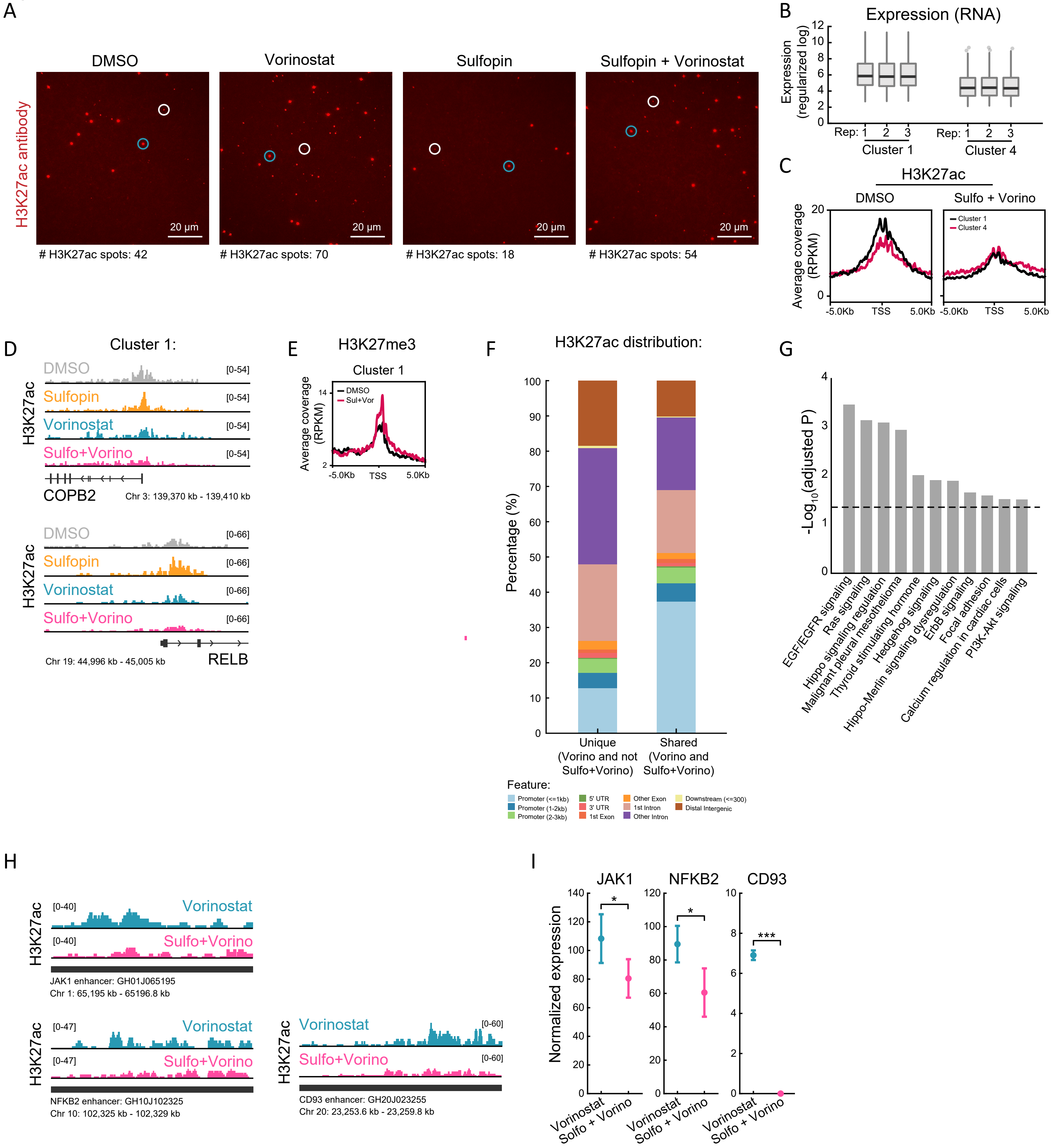
A
DMSO
Vorinostat
Sulfopin
Sulfopin + Vorinostat
H3K27ac antibody
20 µm
### H3K27ac spots: 42
### H3K27ac spots: 70
### H3K27ac spots: 18
### H3K27ac spots: 54
B
Expression (RNA)
Expression (regularized log)
0
2
4
6
8
10
12
Rep: 1 2 3
Cluster 1
Rep: 1 2 3
Cluster 4
C
H3K27ac
DMSO
Sulfo + Vorino
Average coverage (RPKM)
20
0
Cluster 1
Cluster 4
-5.0Kb
TSS
5.0Kb
D
Cluster 1:
H3K27ac
DMSO
Sulfopin
Vorinostat
Sulfo+Vorino
[0-54]
COPB2
Chr 3: 139,370 kb - 139,410 kb
[0-66]
Chr 19: 44,996 kb - 45,005 kb
RELB
E
H3K27me3
Cluster 1
Average coverage (RPKM)
14
2
DMSO
Sul+Vor
-5.0Kb
TSS
5.0Kb
F
H3K27ac distribution:
Percentage (%)
0
10
20
30
40
50
60
70
80
90
100
Unique (Vorino and not Sulfo+Vorino)
Shared (Vorino and Sulfo+Vorino)
Feature:
Promoter (<=1kb)
Promoter (1-2kb)
Promoter (2-3kb)
5' UTR
3' UTR
1st Exon
Other Exon
1st Intron
Other Intron
Downstream (<=300)
Distal Intergenic
G
$-Log_{10}$(adjusted P)
0
1
2
3
4
EGF/EGFR signaling
Ras signaling
Hippo signaling regulation
Malignant pleural mesothelioma
Thyroid stimulating hormone
Hedgehog signaling
Hippo-Merlin signaling dysregulation
ErbB signaling
Focal adhesion
Calcium regulation in cardiac cells
PI3K-Akt signaling
H
H3K27ac
[0-40]
Vorinostat
Sulfo+Vorino
JAK1 enhancer: GH01J065195
Chr 1: 65,195 kb - 65196.8 kb
[0-47]
NFKB2 enhancer: GH10J102325
Chr 10: 102,325 kb - 102,329 kb
[0-60]
CD93 enhancer: GH20J023255
Chr 20: 23,253.6 kb - 23,259.8 kb
I
JAK1
NFKB2
CD93
Normalized expression
0
20
40
60
80
100
120
140
*
***
2
4
6
8
10
Vorinostat
Solfo + Vorino

**Figure S3: H3K27ac levels decrease following the combined treatment specifically on oncogenes**

(A) Representative single-molecule images of individual H3K27ac nucleosomes imaged by TIRF microscopy (as described in Fig. 3A), in SU-DIPG13 treated as described in Fig. 3B. White circles indicate antibody spots, and blue circles indicate TetraSpeck beads that are used for the alignment with the nucleosomes. Number of H3K27ac antibody spots is indicated in each FOV.

(B) Averaged expression levels of genes comprising cluster 1 and cluster 4, in control DMSO-treated SU-DIPG13 cells. Three technical replicates are shown. Cluster 1 genes show higher expression levels under basal conditions compared to clusters 4 genes.

(C) H3K27ac Cut&Run averaged signal over the TSS of genes comprising cluster 1 and cluster 4, in SU-DIPG13 cells treated as described in Fig. 3D.

(D) IGV tracks of the cluster 1' genes- *RELB* and *COPB2*, showing H3K27ac coverage upon their promoters in SU-DIPG13 treated as described in Fig. 3D.

(E) H3K27me3 Cut&Run averaged signal over the TSS of genes comprising cluster 1.

(F) Proportion of H3K27ac peaks corresponding to the indicated genomic features. Shown are peaks that were detected both in Vorinostat and Sulfopin+Vorinostat treated cells (shared peaks, bottom line), as well as H3K27ac peaks that were only present in Vorinostat-treated cells and were lost in the combined treatment with Sulfopin (unique peaks). H3K27ac peaks that were lost in the combined treatment with Sulfopin are associated with distal genomic regions.

(G) Functional enrichment analysis of the genes linked to enhancers (top targets of high confident enhancers), marked with H3K27ac exclusively in SU-DIPG13 cells treated with Vorinostat, and are lost in the combined treatment. gProfiler algorithm[126] was used to calculate enrichment against the WikiPathways genesets[129]. Dashed line denotes adjusted p-value = 0.05. Genes associated with Vorinostat-unique enhancers are enriched for oncogenic signaling pathways.

(H) IGV track of *JAK1, NFKB2* and *CD93* enhancer locus, showing H3K27ac coverage in SU-DIPG13 treated with 1uM Vorinostat or the combination of 10uM Sulfopin and 1uM Vorinostat.

(I) Normalized expression levels of *JAK1, NFKB2* and *CD93* genes in SU-DIPG13 treated as in F. Mean +/- SD of three technical repeats is shown. *P < 0.05; ***P<0.001 (two-sample t-test).

Figure S4

A

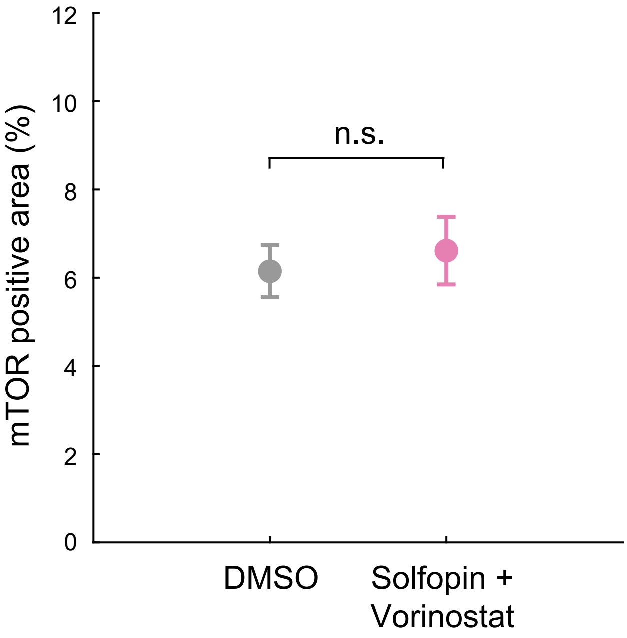

12
10
8
6
4
2
0
mTOR positive area (%)
n.s.
DMSO
Solfopin +
Vorinostat


B

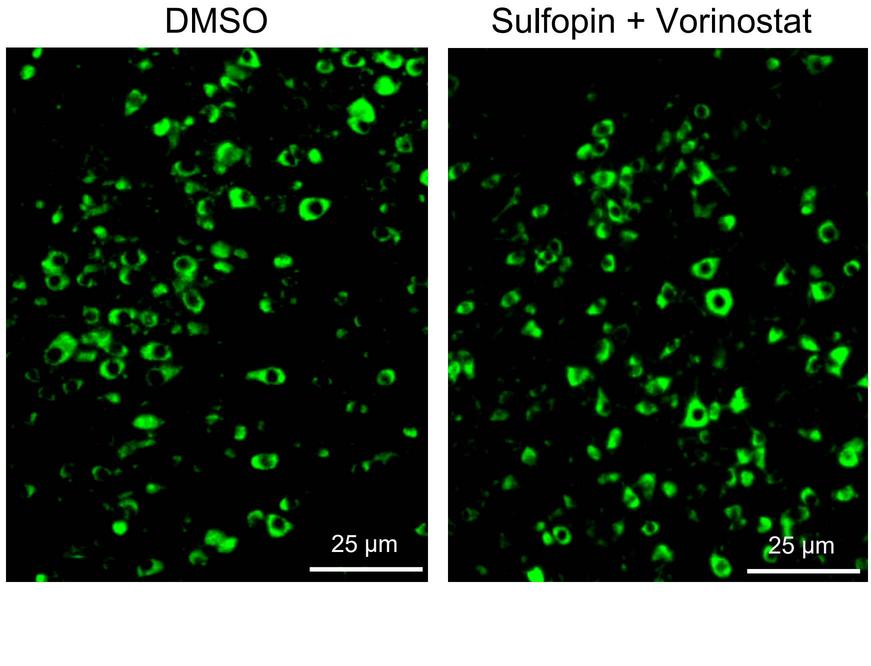

DMSO
Sulfopin + Vorinostat
25 μm
25 μm

**Figure S4: The combined treatment with Sulfopin and Vorinostat results in tumor growth reduction *in-vivo***

(A-B) mTOR immunofluorescent signal (% of positive area) detected in ROI of H3-K27M negative cells (mouse cells), from brain sections of mice treated with either DMSO or the combination of Sulfopin and Vorinostat. N.S. indicates $P > 0.05$ (two-sample t-test). Representative images are shown in B.

Table S1-Antibodies used throughout the study

| **REAGENT or RESOURCE** | **SOURCE** | **IDENTIFIER** |
|---|---|---|
| Histone H3 (K27M Mutant Specific) (D3B5T) Rabbit mAb | Cell signaling | Cat#74829; RRID:AB_2799861 |
| Peroxidase-AffiniPure Goat Anti-Rabbit IgG | Jacksoon Immuno Research Laboratory | Cat#111-035-144; RRID:AB_2307391 |
| Acetyl-Histone H3 (Lys27) (D5E4) XP® Rabbit mAb | Cell signaling | Cat#8173; RRID:AB_10949503 |
| Acetyl-Histone H3 (Lys27) (D5E4) XP(R) Rabbit mAb (Alexa Fluor 647 Conjugate) | Cell signaling | Cat#39030; RRID:AB_2799145 |
| Tri-Methyl-Histone H3(K27) (C36B11) Rabbit mAb | Cell signaling | Cat#9733; RRID:AB_2616029 |
| Histone H3 (D1H2) XP® Rabbit mAb | Cell signaling | Cat#4499; RRID:AB_10544537 |
| mTOR (7C10) Rabbit mAb | Cell signaling | Cat#2983; RRID:AB_2105622 |
| p21 Waf1/Cip1 (12D1) Rabbit mAb | Cell signaling | Cat#2947; RRID: AB_823586 |
| Phospho-S6 Ribosomal Protein (Ser235/236) (D57.2.2E) XP® Rabbit mAb | Cell signaling | Cat#4858; RRID:AB_916156 |
| Phospho-mTOR (Ser2448) Antibody | Cell signaling | Cat#2971; RRID:AB_330970 |
| Recombinant Anti-ATF3 antibody [EPR19488] - ChIP Grade | Abcam | Cat#207434; RRID:AB_2734728 |
| Monoclonal Anti-β-Tubulin I | Sigma | Cat#T7816-100UL; RRID:AB_261770 |

Table S2-Oligonucleotides used throughout the study

| **Gene** | **FWD/REV** | **Sequence** |
|---|---|---|
| Pes1 | FWD | TCTTCCTGTCCATCAAAGGC |
| Pes1 | REV | GTGGCCATGACCCTGTAGTC |
| HSP90AB1 | FWD | CCAGGCACTTCGGGACAACTC |
| HSP90AB1 | REV | CAAGGGAAAAGCCAGAAGATAGCA |
| NCL | FWD | ACTGACCGGGAAACTGGGTC |
| NCL | REV | TGGCCCAGTCCAAGGTAACT |
| E2F2 | FWD | ACAAGGCCAACAAGAGGCTG |
| E2F2 | REV | TCAGTCCTGTCGGGCACTTC |
| RNA5S1 | FWD | GGCCATACCACCCTGAACGC |
| RNA5S1 | REV | CAGCACCCGGTATTCCCAGG |
| RNA5-8SN4 | FWD | GCTCTAACCTTACCTACCTGG |
| RNA5-8SN4 | REV | TGAGCCATTCGCAGTTTCAC |
| MYC | FWD | CAGCTGCTTAGACGCTGGATT |
| MYC | REV | GTAGAAATACGGCTGCACCGA |
| mTOR | FWD | TATCCGCTACTGTGTCTTGGC |
| mTOR | REV | CTCTGTCAGGATCTGGATGAGC |
| PSMD3 | FWD | GAGTTCCTGGACAAGCTGGA |
| PSMD3 | REV | AGGTAATTCCGCAGCAGGAG |
| NOC4L | FWD | GCCACCCCTCCTTTCAGG |
| NOC4L | REV | GGGGAACAGGTAGTTGCCTT |
| GAPDH | FWD | GGTGTGAACCATGAGAAGTATGA |
| GAPDH | REV | GAGTCCTTCCACGATACCAAAG |

Table S3-Drugs and inhibitors used throughout the study

| REAGENT or RESOURCE | SOURCE | IDENTIFIER |
|---|---|---|
| Sulfopin | Gift from Dr. Nir London’s Lab | |
| Vorinostat | MCE | Cat#HY-10221 |
| Vorinostat | LC | Cat#V-8477 |
| MM-102 | Selleck | Cat#S7265 |
| EPZ6438 | Selleck | Cat#S7128 |
| GSK-J4 | Selleck | Cat#S7070 |
